# Zebrafish *adamtsl4* knockout recapitulates key features of human *ADAMTSL4*-related diseases: a gene involved in extracellular matrix organization, cell junctions and development

**DOI:** 10.1101/2025.01.16.632701

**Authors:** Angel Tevar, Jose-Daniel Aroca-Aguilar, Raquel Atiénzar-Aroca, Ana I. Ramírez, José A. Fernández-Albarral, Julio Escribano

## Abstract

Loss-of-function mutations in *ADAMTSL4,* a gene encoding an extracellular matrix-associated protein with incompletely understood biological roles, are linked to autosomal recessive disorders predominantly characterized by lens dislocation. Pupil ectopia, increased intraocular pressure, retinal detachment, cataracts, and skeletal abnormalities are also observed in some patients. To investigate *ADAMTSL4* biology and related diseases we established a zebrafish knockout line using CRISPR/Cas9 genome editing. The generated zebrafish model harboured the c.234-351del mutation in *adamstsl4*, reducing its mRNA levels by 75% in 6 days post fertilization (dpf) larvae, and predicting to produce an inactive protein (p.(Gln78Hisfs*127)). Forty percent of F3 knockout larvae (6 dpf) displayed lethal phenotypes characterized by multiple ocular and non-ocular developmental defects, including pericardial, perivitelline and periocular edema, absence of swim bladder, craniofacial malformations and microphthalmia. The remaining 60% larvae survived and displayed only reduced pupil area, indicating incomplete penetrance of the lethality. Histology revealed extracellular matrix (ECM) and intercellular junctions abnormalities within the cornea, iris, lens, and retinal pigment epithelium (RPE). Adult knockout zebrafish (6 months) presented phenotypes resembling ectopia lentis et pupillae and craniosynostosis, with optical defects in the lens and impaired visual function. ECM and cell junction disorganization in the cornea, lens and RPE were also present in these animals. Transcriptomic analysis revealed disrupted expression of genes involved in development, ECM and cell junctions among other biological processes. These findings show that a*damtsl4* recapitulates key features of human *ADAMTSL4*-related disorders and that this gene is essential for normal ECM structure, cell junctions and embryonic development.

**Highlights:**

- Zebrafish *adamtsl4* knockout recapitulates human *ADAMTSL4*-related phenotypes.
- *adamtsl4* is involved in embryonic development.
- *adamtsl4* participates in ECM organization and cell adhesion.

## 1. Introduction

Functional alteration of *ADAMTSL4* (ADAM metallopeptidase with thrombospondin type 1 motif-like 4) underlie a spectrum of autosomal recessive diseases, ranging from the rare combination of cranial and ocular alterations in the case of craniosynostosis with ectopia lentis [1, 2] to restricted ocular alterations, such as autosomal recessive isolated ectopia lentis [3, 4] and ectopia lentis et pupillae [5]. Additional minor eye anomalies with no displacement of the pupil and very mild displacement of the lens are associated with functional disruption of *ADAMTSL4*, including congenital abnormalities of the iris, refractive errors that may lead to amblyopia and early-onset cataract [6], childhood glaucoma [7], increased intraocular pressure and retinal detachment and increased ocular axial length [3, 8]. Notably, the presentation of *ADAMTSL4*-related ocular abnormalities can vary both among affected individuals within the same family and between the eyes of a single individual, underscoring the influence of modifier genes on the phenotypes. Recently, it has also been reported that partial functional defects in *ADAMTSL4* may contribute to increased susceptibility to early-onset glaucoma [9]. ADAMTSL4 is a member of the ADAMTS (a disintegrin and metalloproteinase with thrombospondin motifs) family of extracellular matrix (ECM) proteins. These proteins are involved in various physiological processes, including tissue development, homeostasis, and ECM remodeling. ADAMTS proteins comprise a protease domain and an ancillary domain that is involved in substrate recognition and in tissue specificity [10]. ADAMTS-like proteins are structurally related to ADAMTS proteases but lack the characteristic ADAMTS protease domains, making it unlikely that they exhibit protease activity. The biological functions of ADAMTSL proteins are not yet fully understood. However, evidences suggest they play important roles in extracellular matrix ECM organization, connective tissue maintenance, the regulation of fibrillin microfibrils [11, 12] and growth factor signalling [13, 14]. The ADAMTSL4 protein contains several functional domains essential for its role in maintaining the extracellular matrix. It contains seven thrombospondin type 1 repeats (TSR1) which facilitate binding to ECM proteins, with most of them clustered in the C-terminal region of the protein. Additionally, spacer domains are present, which mediate protein-protein interactions [10]. Integrity of the functional domains is required for the structural and cellular signaling roles within connective tissues, contributing to processes such as craniofacial development and ocular function.

Most *ADAMTSL4* pathogenic mutations appear to lead to protein truncation either disrupting these domains/or leading to NMD mRNA degradation. The precise mechanisms by which *ADAMTSL4* mutations contribute to the development of different pathological phenotypes remain largely unexplored and unclear. Functional disruption of this gene in mice originates focal retinal pigment epithelium (RPE) defects and mimics the human ectopia lentis phenotype, suggesting that it is required for stable anchorage of zonule fibers to the lens capsule [15]. In this study, we aimed to generate a zebrafish line knockout gene to establish a novel animal model for studying *ADAMTSL4* biology and related diseases. Our results show that this gene is essential for extracellular matrix structure and cell junctions, playing a role in embryonic, cranial and ocular development in zebrafish and that its inactivation in this animal mimics key ocular and non-ocular features of *ADAMTSL4*-related disorders.

## 2. Materials and Methods

### 2.1. Animals

Wild-type AB zebrafish (Danio rerio) were kept in a controlled environment with a 14-hour light cycle followed by a 10-hour dark cycle at a 28 °C temperature. They were provided with a standard diet based on established protocols [16]. Zebrafish embryos were raised in E3 medium at a temperature of 28 °C. The E3 medium had a pH of 7.2 and consisted of 5 mM NaCl, 0.17 mM KCl, 0.33 mM CaCl2, 0.33 mM MgSO4, and 0.0001% methylene blue. To facilitate imaging on a Nikon SMZ18 mircoscope and a DS-Ri2 camera, zebrafish embryos and larvae were anesthetized using 0.02% tricaine methanesulfonate (MS222, Sigma-Aldrich) and immobilized using a 2% methylcellulose solution. Adult fishes were anesthetized with 0.04% tricaine methanesulfonate and euthanized by prolonged immersion in MS222 (200–300 mg/L). All animal care and experiments adhered to the guidelines set by the Institutional Animal Research Committee of the University of Castilla-La Mancha and were approved under the reference number PR-2017-01-19.

### 2.2. ADAMTSL4 gene homology analysis

The human and zebrafish *ADAMTSL4* gene homology was evaluated using the Ensembl database (https://www.ensembl.org/index.html). Gene structure and sinteny were analyzed using the *Ensembl region comparison tool*. The Basic Local Alignment Search Tool (BLAST) (https://blast.ncbi.nlm.nih.gov/Blast.cgi) was used to determine the percentage of amino acid sequence identity. The three-dimensional structures of the orthologous proteins were obtained via structural modeling using deep learning and artificial intelligence through the AlphaFold Protein Structure Database server (https://alphafold.ebi.ac.uk/), and they were superimposed using the FATCAT server [17]. The files provided by these servers in PDB format were visualized using the Jmol 3D structure viewer. Tertiary structures of orthologous proteins with superposition values of P<0.05 were considered significantly similar.

### 2.3 CRISPR/Cas9 gene editing

We used a two-part guide RNA system (tracrRNA/crRNA). Target selection and the design of crRNA were carried out using a customized Alt-R^TM^ CRISPR-Cas9 guide RNA from Integrated DNA Technologies (IDT, Coralville, IA, USA) (https://eu.idtdna.com/site/order/designtool/index/CRISPR_CUSTOM). The assessment of potential off-target sites and the determination of crRNA’s highest on-target activity were performed using the IDT CRISPR-Cas9 guide RNA design checker (https://eu.idtdna.com/site/order/designtool/index/CRISPR_SEQUENCE). Two crRNAs targeting exon 3 of adamtsl4 were designed (adamtsl4E3g1: 5’-GATGGTCTATTCCATGTAAGGTTTTAGAGCTATGCT-3’ and adamtsl4E3g2: 5’-TATCCTCCTAATGAACCATAGTTTTAGAGCTATGCT-3’). To generate deletions of approximately 100 bp in the exon 3 of *adamtsl4*, the two crRNAs were microinjected simultaneously. To that end, the two crRNAs (36 ng/µl each) were mixed and incubated with the tracrRNA (67 ng/µl) at 95 °C for 5 min and cooled to room temperature to allow hybridization. Then, the Alt-R® CRISPR-Cas9 protein (250 ng/µl) was mixed with the tracrRNA/crRNA complex and incubated for 10 min at 37 °C to form the ribonucleoprotein (RNP) complex. Approximately 3 nl of the RNP complex were injected into the animal pole of one-cell stage embryos (50-250 embryos per experiment). The injection procedure was performed using a Femtojet 5247 microinjector (Eppendorf, Hamburg, Germany) under a Nikon SMZ18 stereomicroscope. As a negative control, embryos were injected with Cas9/tracrRNA in the absence of crRNA. All reagents for CRISPR/Cas9 gene editing were provided by IDT.

### 2.4. Genotyping of CRISPR/Cas9-Induced Mutations by PCR and Sanger Sequencing

PCR-ready genomic DNA was extracted from entire zebrafish embryos at 5 days post-fertilization (dpf) as well as from the caudal fin of anesthetized adult zebrafish employing the HotSHOT method [18]. Tissue samples were incubated with 20 µL base solution (25 mM KOH, 0.2 mM EDTA) at 95 °C for 30 min followed by the addition of 20 µL neutralization buffer (40 mM TrisHCl, pH 5). Mutations in *adamtsl4* exon 3 were analyzed by PCR using the primers (adamtsl4E3Fw1: 5’-CAGGCAGCAGGACGGCAG-3’ and adamtsl4E3Rv1: 5’-CCATGACTCGGTGTCCGTGC-3’) in the following conditions: an initial denaturation step at 95 °C for 3 min followed by 35 cycles consisting of denaturation at 95° C for 30 s, annealing at 63 °C for 30 s and extension at 72° C for 45 s. A final extension step at 72° C for 5 min was also included. PCR products were analyzed either by 1% agarose gel electrophoresis or direct Sanger sequencing (STAB VIDA, Caparica, Portugal).

### 2.5. Morphological characterization of zebrafish

Larvae (6 dpf) and adult zebrafish (5 mpf) were photographed from lateral, ventral, and dorsal views using a Nikon SMZ18 stereo zoom microscope. The volumes of the anterior chamber and the lens within the anterior chamber were estimated under the assumption that they are spherical caps. These volumes were calculated using the formulas:

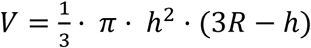

and

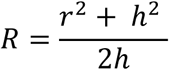

where R is the radius of the sphere, h is the height of the cap and r is the radius of the cap. The areas of the lens nuclei were quantified using the NIS-Elements software (Nikon). At least three lenses were analyzed per genotype.

### 2.6. Dissection of Adult Zebrafish Lens

Adult zebrafish were euthanized following institutional ethical guidelines using a 0.02% tricaine methane sulfonate (MS-222) solution. Eyes were extracted and transferred to a petri dish containing phosphate-buffered saline (PBS) at room temperature. The lens was carefully removed and immediately transferred to fresh PBS. The optical properties of the lens were assessed by analyzing the transmission and reflection of light under vertical and horizontal illumination sources.

### 2.7. Light microscopy

Zebrafish larvae (6 dpf) were fixed in a solution containing 2.5% glutaraldehyde and 4% paraformaldehyde in 0.1 M phosphate buffer Milloning (PBM, pH 7.4) for 4 h at 4°C and postfixed in 1% osmium tetroxide for 1.5 h at 4°C. Tissue samples were dehydrated using series of increasing concentrations of acetone (30-100%) and embedded in araldite. Semi-thin sections (0.5 μm) were obtained using a ultramicrome Leica Ultracut UCT, stained with Toluidine blue and analyzed by light microscopy. Adult zebrafish heads (5 months) were fixed overnight in 4% paraformaldehyde (4% PFA and 0.1 M PBS, pH 7.3), cryoprotected by immersion for two days at 4°C in a solution of 30% sucrose in 0.1 M PBS, embedded in 10% porcine gelatin and 15% sucrose, and stored at -80°C. Serial cryosections were obtained using a Leica CM3050 S cryostat (14 μm). At least three larvae or adult zebrafish heads were used per genotype. For histological staining, cryosections were incubated with Harris hematoxylin solution (HHS80, Sigma-Aldrich) and an alcoholic eosin solution (HT1101116-500ML, Sigma-Aldrich) following standard protocols. The slides were then prepared by mounting medium using Cytoseal (8311-4, Thermo Scientific, Waltham, MA, USA). Craniofacial defects were analyzed by whole-mount bone and cartilage staining, using a cartilage staining solution (0.02% Alcian blue, 05500-5G, Sigma-Aldrich) followed by a bone staining solution (0.05% alizarin red S, A5533-25G, Sigma-Aldrich)[19].

### 2.8. Transmission electron microscopy (TEM)

The tissue samples were promptly fixed in a solution of 2.5% glutaraldehyde and 4% paraformaldehyde in 0.1 M phosphate buffer Milloning (PBM, pH 7.4) for 4 h at 4 °C. After washing in PBM, the samples were postfixed in 1% osmium tetroxide for 1.5 h at 4 °C. Following additional washing steps in PBM, the tissues were dehydrated in an ascending series of acetone (30-100%) and then embedded in araldite. The araldite blocks were sectioned. Initially, semi-thin sections of 0.5 μm thickness were stained with Toluidine blue and examined under a light microscope to identify the optimal regions for TEM. Subsequently, the selected araldite blocks were trimmed and ultramicrotomy was performed using a Reichert OM-V3 ultramicrotome (Wetzlar, Germany). Thin sections were stained with uranyl acetate and lead citrate for contrast. Finally, these sections were analyzed using a JEOL JEM 1010 transmission electron microscope.

### 2.9. Optokinetic Response in Adult Zebrafish

Adult zebrafish (11 months) visual function was measured using optokinetic response (OKR) through smooth-pursuit and rapid saccade eye movements measurement [20, 21]. The experiments were conducted in the morning under constant illumination and water temperature (27 °C). Four wild-types and four *adamtsl4* KO zebrafish were analyzed.

### 2.10 Transcriptome Analysis by RNAseq

RNA was extracted from homozygous *adamtsl4* loss-of-function pools of 50 larvae (6 dpf) and pools of 4 adult zebrafish eyes (5 months). The TRI reagent (SIGMA) was used for isolation following the manufacturer’s instructions. Wild-type zebrafish siblings were used as control. The isolated RNA was further purified using RNAeasy columns (Quiagen, Germantown, MD, USA) before being sent to Macrogen Next-Generation Sequencing Division (Macrogen, Inc., Seoul, Korea) for high-throughput sequencing. Two independent biological replicates were analyzed per genotype. Libraries were prepared using the TruSeq Stranded mRNA LT Sample Prep Kit (Illumina, Foster City, CA, USA), which allows for the capture of coding RNA and various forms of polyadenylated non-coding RNA. Paired-end sequencing with a read length of 150 bp was performed using the NovaSeq6000 System (Illumina). Trimmomatic 0.38 (Bolger et al., 2014) was used to remove low-quality bases and adapter sequences from the reads. The reads were then aligned to the zebrafish genome reference (GRCz11) using the HISAT2 aligner (Kim et al., 2015). Expression profiles were determined based on read count and a normalization value that accounted for transcript length and coverage depth. Differential expression analysis was conducted to compare KO *adamtsl4* to wild-type zebrafish. Reads per kilobase of transcript per million mapped reads (RPKM) were used for this analysis. Genes with a fold change of 2.0 or greater and a p-value less than 0.05 between the different biological replicates were considered differentially expressed genes (DEGs). ShinyGO pathway enrichment analysis was employed to explore the functional significance of the DEGs. Biological pathways were ranked based on their false discovery rate (FDR) [22] and the 20 most significant biological pathways were selected for further analysis.

### 2.11. Quantitative Reverse Transcription PCR (qRT-PCR)

RNA was isolated from pools of 50 zebrafish larvae (6 dpf) and pools of 4 adult zebrafish eyes (5 months) using the RNeasy Minikit (#74104, Qiagen, Germantown, MD, USA). The RNA samples were treated with RNase-free DNase I following the instructions provided by the manufacturer. The purified RNA was used to synthesize cDNA using the RevertAid First Strand cDNA Synthesis Kits (#K1622, Thermo Fisher Scientific, Waltham, MA, USA). The expression levels of adamtsl4 mRNA or specific DEGs relative to ef1α mRNA were determined using the 2−ΔΔCt method [23], employing the primer pairs described in Supplementary Table S1. For the PCR analysis, 1 μL of cDNA was used as a template in a reaction volume of 10 μL, which included 5 μL of Power SYBR Green PCR Master Mix (Thermo-Fisher Scientific) and 200 nM of each primer. The thermocycling conditions consisted of an initial denaturation step at 95°C for 10 minutes, followed by 40 cycles of denaturation at 95°C for 15 seconds, annealing and extension at 60°C for 40 seconds. The PCR products and their dissociation curves were analyzed using a 7500 Fast real-time PCR system thermal cycler (Thermo-Fisher Scientific). A qRT-PCR negative control was included for each specific primers pair by omitting the template cDNA. The mean expression values for each sample were calculated using qRT-PCR results obtained from three independent experiments.

### 2.12. Statistics

Student’s t-test was employed for evaluation of statistical comparisons between groups using the SigmaPlot 12.0 software (Systat Software Inc., San Jose, CA, USA).

## 3. Results

### 3.1. Evolutionary conservation analysis

To assess the feasibility of using zebrafish as an animal model to study the role of *ADAMTSL4* in human ocular diseases we analyzed its evolutionary conservation. The zebrafish *adamtsl4* gene (LO018340.1) is located on chromosome 16 while the human gene is localized in the long arm of chromosome 1. The synteny analysis revealed conservation of the gene order in the regions immediately flanking this gene in the human and zebrafish genomes (Supplementary Fig. S1A). Comparison of the structure of the human and zebrafish genes showed 17 coding exons in both genes, with a high degree of conservation in the sequence of 10 of them (exons 5 to 8 and 12 to 17) and in the intron-exon organization (Supplementary Fig. S1B). The amino acid sequence identity between the two ortholog proteins was 39.58%, with the highest degree of conservation located in the C-terminal region of the two proteins (Supplementary Fig. S2A). The analysis of the three-dimensional conservation by homology-based structural modeling showed that the predicted structures were significantly similar (p = 8.40 x 10^-4^) and overlapped quite well (Supplementary Fig. S2B). The evolutionary conservation observed between the two ortholog genes supports zebrafish as a suitable animal model for studying *adamtsl4*-associated pathologies.

### 3.2. Generation of a Zebrafish Line Knockout (KO) for the adamtsl4 Gene Using CRISPR/Cas9 Genome Editing

To create the zebrafish *adamtsl4* KO line, we designed two specific crRNAs targeting exon 3 of the gene. The two crRNAs were co-injected simultaneously into zebrafish embryos at the one-cell stage (Supplementary Fig. S3A), as described in Materials and Methods, to generate deletions of approximately 100 bp, which are easily identifiable by PCR. Following injection, 10 F0 embryos were grown into adult zebrafish and screened for germline-transmitted *adamstsl4* deletions by PCR of exon 3 and electrophoretic analysis (Supplementary Fig. S3B), as indicated in the Materials and Methods section. We selected as the founder an F0 zebrafish that transmitted a deletion of approximately 100 bp. Possible off-target mutations were segregated by two generations of outbreeding (Supplementary Fig. S3A). F2 mutants were inbred to obtain F3 mutant homozygotes, which were identified by PCR analysis. Characterization of this mutation by Sanger sequencing revealed a 118-bp deletion c.234_351del (Fig. 1A). This mutation is predicted to generate a frameshift and a premature termination codon (p.(Gln78Hisfs*127)), that may result in *adamtsl4* mRNA degradation due to the nonsense-mediated mRNA decay (NMD) mechanism. The *adamtsl4* mRNA levels associated with the three genotypes of the offspring produced by inbreeding F2 heterozygotes were analyzed using RT-qPCR. cDNA was synthesized from pooled samples of 50 larvae per genotype. A genotype-dependent reduction in mRNA levels was observed, with heterozygotes showing an approximate 50% reduction and homozygotes around 75%, compared to wild-type levels (Fig. 1B). These results are consistent with the predicted mRNA degradation induced by the NMD mechanism.

**Fig. 1.**
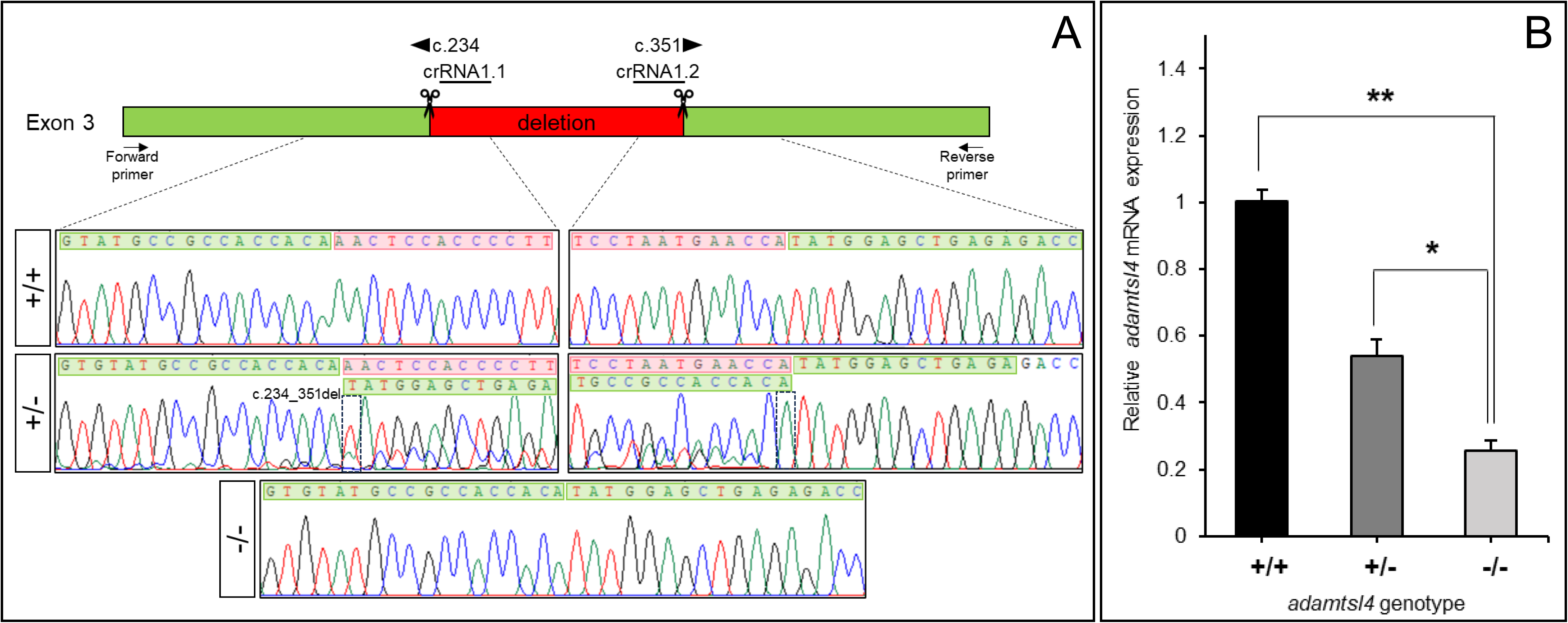
Molecular characterization of the *adamtsl4*-KO zebrafish line generated using CRISPR/Cas9 genome editing. A. Localization of the deletion identified in *adamtsl4* exon 3. Scissors: DNA cleavage sites targeted by two crRNAs (crRNA1.1 and crRNA1.2). The numbers indicate the cDNA nucleotide positions corresponding to the start and end of the deleted DNA fragment. Wild-type sequences are highlighted with a green background, while deleted sequences are marked with a red background. Arrows: position of the forward and reverse primers used for zebrafish genotyping. Rectangles in the electropherograms mark the 5′ and 3′ ends of the deletion. B. Reduced mRNA levels in *adamtsl4*-KO zebrafish larvae. Relative adamtsl4 mRNA levels were measured in 6 days post-fertilization (dpf) zebrafish larvae using RT-qPCR. Data represent the mean ± standard error, derived from three independent experiments performed in triplicate, using pools of 50 larvae. Statistically significant differences between KO and wild-type zebrafish are denoted by asterisks: *p* < 0.05 (*), *p* < 0.01 (**).

### 3.3 Phenotypic characterization of the zebrafish adamstsl4 KO line

The genotype-phenotype correlation associated with *adamtsl4* LoF was evaluated in 113 F3 larvae (6 dpf), obtained by inbreeding F2 mutant zebrafish. Following PCR-based genotyping, larvae were classified by genotype and subjected to morphological analysis (Fig. 2A-H). Although the proportion of homozygous mutant larvae appeared lower than expected, the observed genotype proportions did not deviate significantly from Mendelian expectations: +/+, 31 larvae (29.8%); +/-, 50 larvae (48.1%); and -/-, 23 larvae (22.1%) (χ² test, p=0.5). *Adamtsl4* mutant heterozygous larvae and 60% of KO larvae were viable (Fig. 2I), showing a wild type-like phenotype (Fig. 2B and C, respectively). However, the remaining 40% of KO larvae exhibited severe developmental abnormalities, including microcephaly, microphthalmia, absence of swim bladder, and periocular, pericardial and perivitelline edemas (Fig. 2D), which were associated with lethality in the days following these observations (Fig. 2I). Detailed evaluation of ocular morphology revealed that the viable (wild type-like) KO larvae manifested ocular anomalies (Fig. 4E-G), characterized by a 20% reduction of the relative pupil area (pupil area/total eye area) compared to wild type and heterozygous siblings (Fig. 2J). These findings highlight a critical role for *adamtsl4* in zebrafish embryonic development, particularly in body and ocular morphogenesis.

**Fig. 2.**
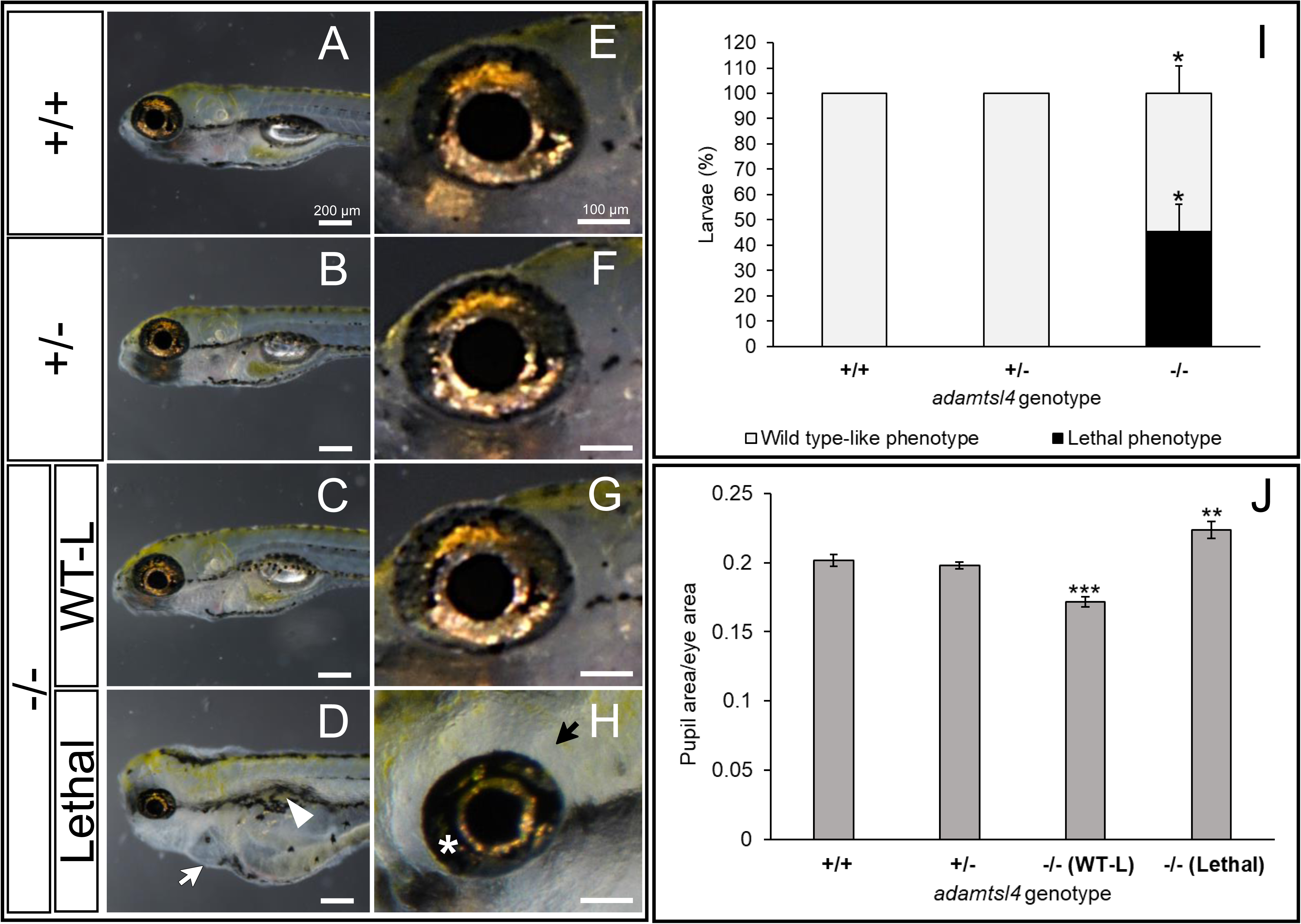
Morphological characterization of F3 *adamtsl4*-KO larvae at 6 dpf. Representative photographs of wild type (+/+), heterozygous (+/-), wild type-like knockout (-/-(WT-L)), and lethal knockout (-/- (Lethal)) larvae are shown. A-D. General body morphology. White arrow: pericardial edema; White arrowhead: abnormal swim bladder. E-H. Magnification of ocular structures. I. Quantification of the percentage of KO larvae displaying wild type-like and lethal phenotypes. The following sample sizes were used: wild type, n = 31; heterozygous, n = 59; homozygous, n = 23. J. Quantification of pupil area normalized to total eye area. The following sample sizes were used for eyes: wild type, n = 24; heterozygous, n = 24; homozygous, n = 24. *: p < 0.05; **: p < 0.01; ***: p < 0.001.

Histological analysis of semi-thin ocular sections using optical microscopy showed that wild type and viable KO larvae showed similar eyeballs size (Fig. 3A and B). In contrast, the eyes of lethal KO larvae were characterized by decreased ocular size and increased surrounding space (Fig. 3C), which is indicative of periocular edema (Fig. 3C, arrows). No apparent differences between ocular anterior segment structures and retina of wild type and KO viable zebrafish larvae were observed (Fig. 3A, B, D, E, G, H, J, K, M and N). However, the lethal KO larvae presented increased foamy appearing pigment cells (xanthophores) particularly evident in the dorsal anterior chamber angle (Fig. 3C, F and I, arrowhead). In addition, the corneal epithelium showed swelling, disrupted cell stratification and intercellular spaces (Fig. 3F-L, asterisks). Intercellular separations were also present between the lens epithelium and lens fiber cells (Fig. 3L, asterisk) and the RPE was enlarged and presented deposits of amorphous material (Fig. 3O, white arrow).

**Fig. 3.**
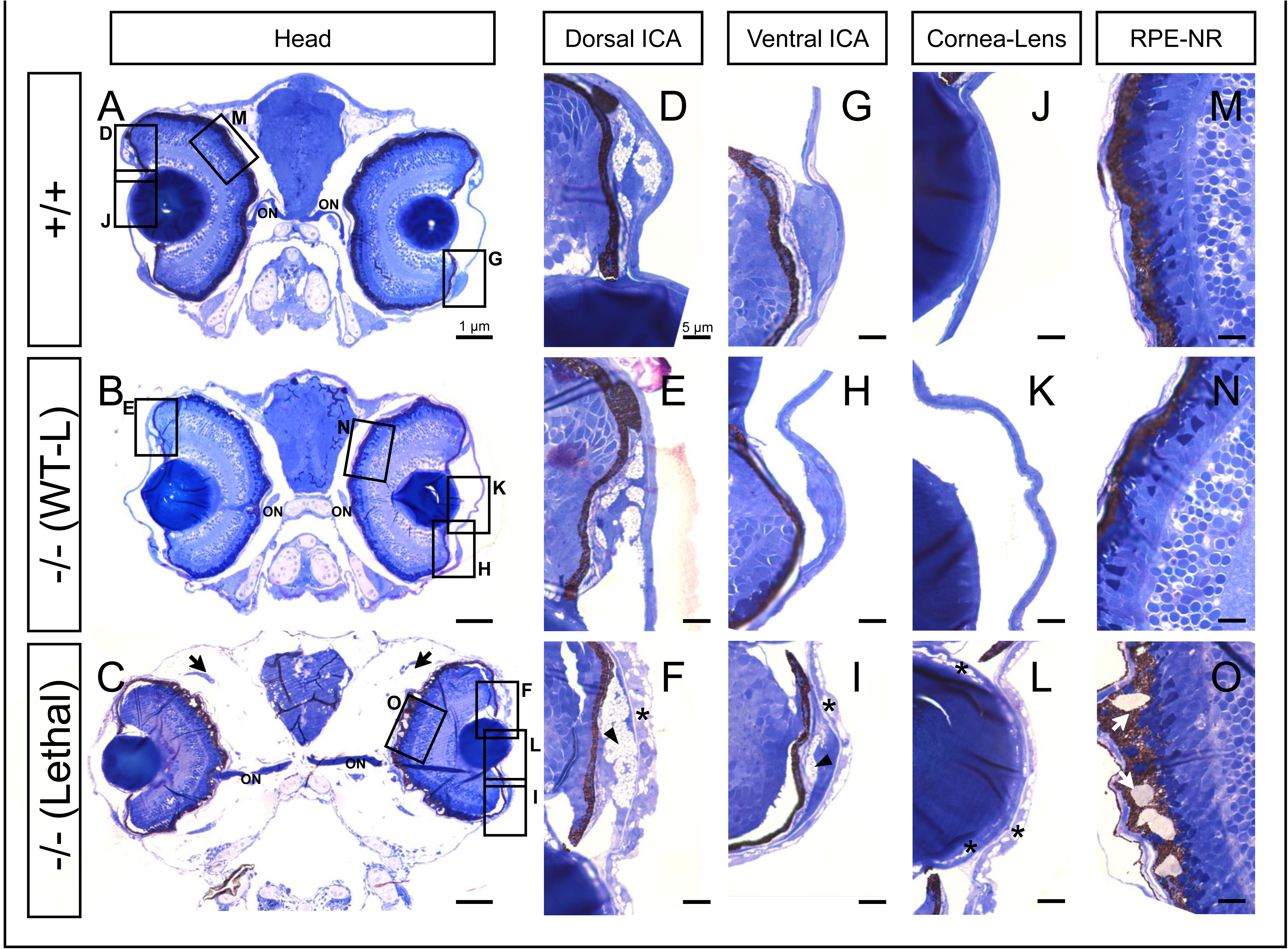
Histological analysis of toluidine blue-stained tissue sections from *adamtsl4* KO zebrafish larvae (6 dpf). Transverse semithin tissue sections (500 nm) were prepared as described in the Materials and Methods section. Representative photographs of wild type (+/+), wild type-like KO (-/- (WT-L)) and lethal KO (-/- (Lethal)) larvae are shown. A-C. Tissue sections from the central part of the eyeball (note the presence of the optic nerve (ON)); rectangles indicate areas magnified in subsequent panels. D-F. Dorsal iridocorneal angle (ICA). G-I. Ventral ICA. J-L. Cornea and lens. M-O. Retinal pigmented epithelium (RPE)-neuroretina (NR) interphase. Asterisks: abnormal intercellular separations; black arrows: periocular edema; black arrowhead: increased xantophores; white arrows: amorphous deposits in the RPE.

Ultrastructural analysis by transmission electron microscopy (TEM) of these specimens confirmed the findings and provided additional information. We observed an increased size of xantophores in both wild-type like and lethal KO larvae (Fig. 4A-C). Additionally, the iridophores presented large guanine crystals compared with wild type larvae (Fig. 4D-F). Swollen cells (indicative of necrosis) and large intercellular separations were evident in the multilayered corneal epithelium of the lethal KO larvae (Fig. 4I, white and black asterisks, respectively) and endothelium (Fig 4I, arrowhead). The wild type corneal stroma presented lamellae formed by abundant and highly ordered and parallel collagen fibrils (Fig 4J, CF), that were progressively less abundant and less structured in the homozygous mutants (Fig. 4K and L, CF). Specifically, the lethal KO larvae exhibited significant disorganization of collagen fibers, with some appearing highly electron-dense (Fig. 4L, arrows). The lens capsule appeared somewhat thickened in both wild type-like and lethal mutant larvae (Fig. 4M-O, white arrow). Interestingly, intercellular spaces were observed between the lens epithelium and lens fibers (Fig. 4O, asterisk), a feature distinct from the autophagy process, seen in the viable KO larvae (Fig. 4N, arrowhead), which is essential for achieving the mature structure of the lens. Additionally, abnormal intercellular spaces were also present between the RPE and neuroretina in the lethal KO larvae (Fig. 4R, asterisks).

**Fig. 4.**
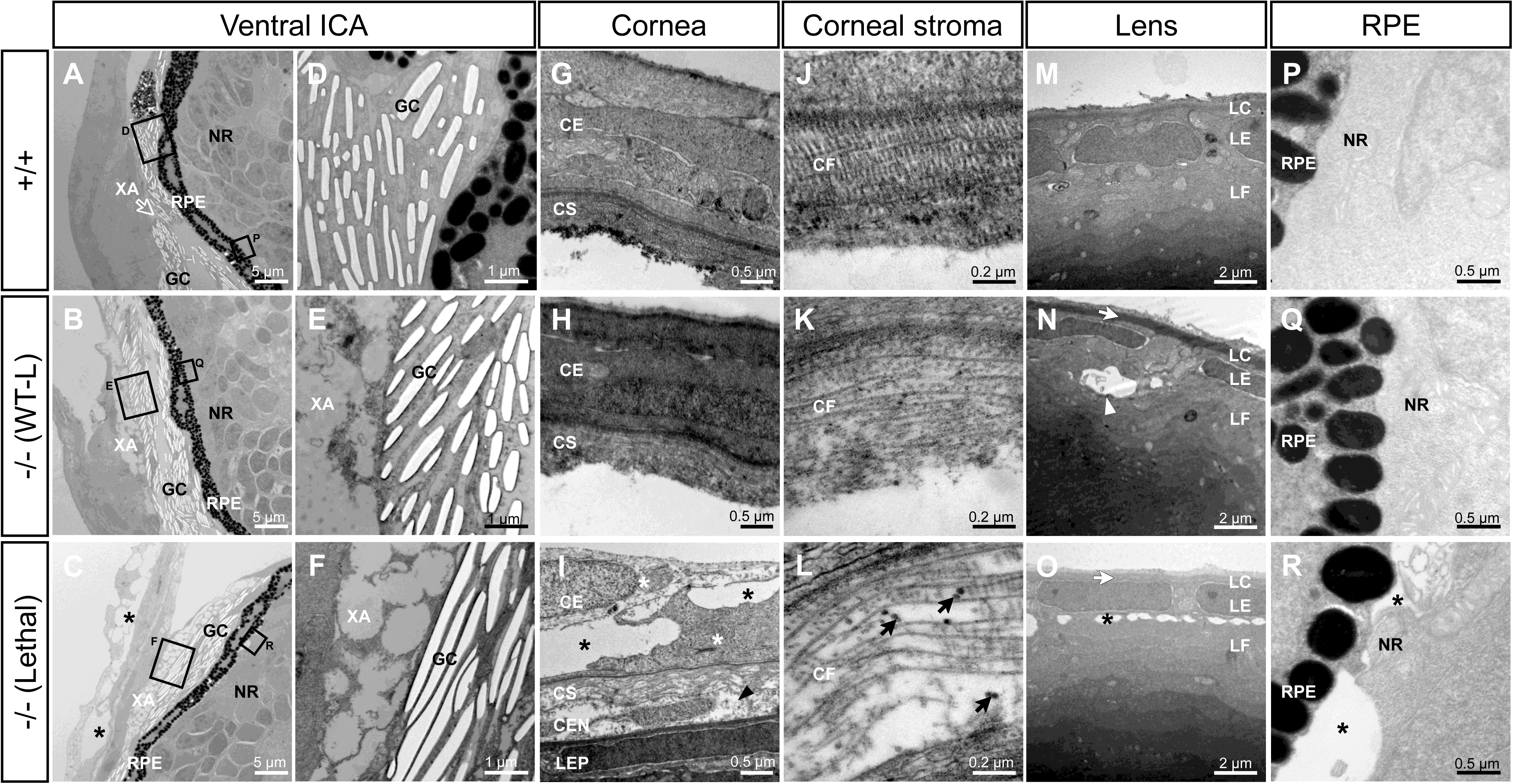
Transmission electron microscopy of the ocular anterior segment in wild type and F3 *adamtsl4* KO zebrafish larvae (6 dpf). Tissue sections were prepared as described in the Materials and Methods section. A-C. Representative photographs of wild type (+/+), wild type-like KO (-/- (WT-L)) and lethal KO (-/- (Lethal)) larvae. ICA: ventral iridocorneal angle. The rectangles indicate the areas magnified in subsequent panels. D-F. Ultrastructural details of iridophores and xantophores (XA), showing the presence of guanine crystals (GC) in the iridophores. G-I. Cornea. Note that the corneal endothelium was lost during tissue preparation in panels G and H. J-L. Ultrastructural details of the corneal stroma. M-O. Lens. P-R. Retinal pigment epithelium (RPE)-neuroretina (NR) interphase. CE, corneal epithelium; CEN, corneal endothelium; CF, collagen fibrils; CS, corneal stroma; GC, guanine crystals; LC, lens capsule; LEP, lens epithelium; LF, lens fibers; NR, neuroretina; RPE, retinal pigment epithelium; XA, xanthophores.. Black arrows: enlarged and electron-dense CF. Black arrowhead: necrotic CE cell; black asterisks: abnormal intercellular separations; white arrowhead: autophagy; white arrows: thickened LC; white asterisks: swollen necrotic cells.

The phenotype of adult F3 zebrafish (5 months old) was also evaluated. KO animals showed altered frontal cranial morphology, which was evident in the lateral view (Fig. 5A and B, arrow) but not observable in the dorsal view (Fig. 5C and D). Bone and cartilage staining using Alcian blue and Alizarin red revealed irregular trajectories in the interfrontal and coronal sutures (Fig. 5E-H), indicative of craniosynostosis. Furthermore, detailed ocular evaluation identified abnormal pupil morphology, marked by ventral elongation, indicative of ectopia pupillae, (Fig. 6A and B, arrow) and a reduction in the relative pupil area by approximately 20% compared to wild type specimens (Fig. 6C). Additionally, an increase in xantophores was noted (Fig. 6B, asterisk). Morphometric analysis revealed significant reduced volumes of both the anterior chamber (Fig. 6D-F) and the portion of the lens within this chamber in the KO zebrafish (Fig. 6D-E and G), that can be the result of ectopia lentis.

**Fig. 5.**
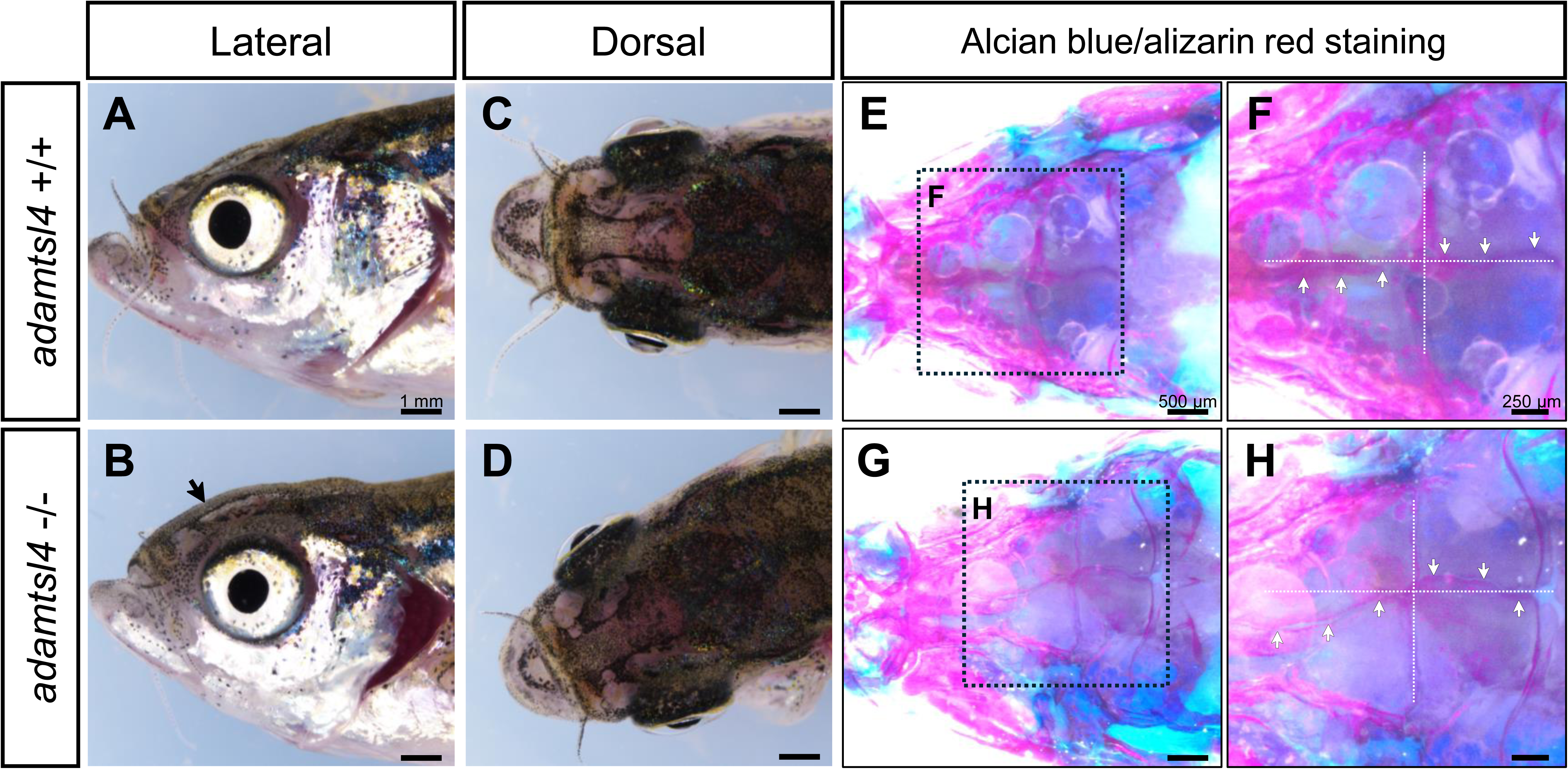
Craniofacial characterization of 5 months F3 *adamtsl4*-KO zebrafish. A-D. Representative images of wild-type and *adamtsl4*-KO zebrafish before clearing and staining, highlighting gross craniofacial morphology. E-H. Staining with alcian blue for cartilage and alizarin red for bone structures reveals detailed craniofacial anatomy. Rectangles indicate regions of interest magnified in subsequent panels, providing a closer examination of structural differences between genotypes. White arrows indicate intracranial sutures.

**Fig. 6.**
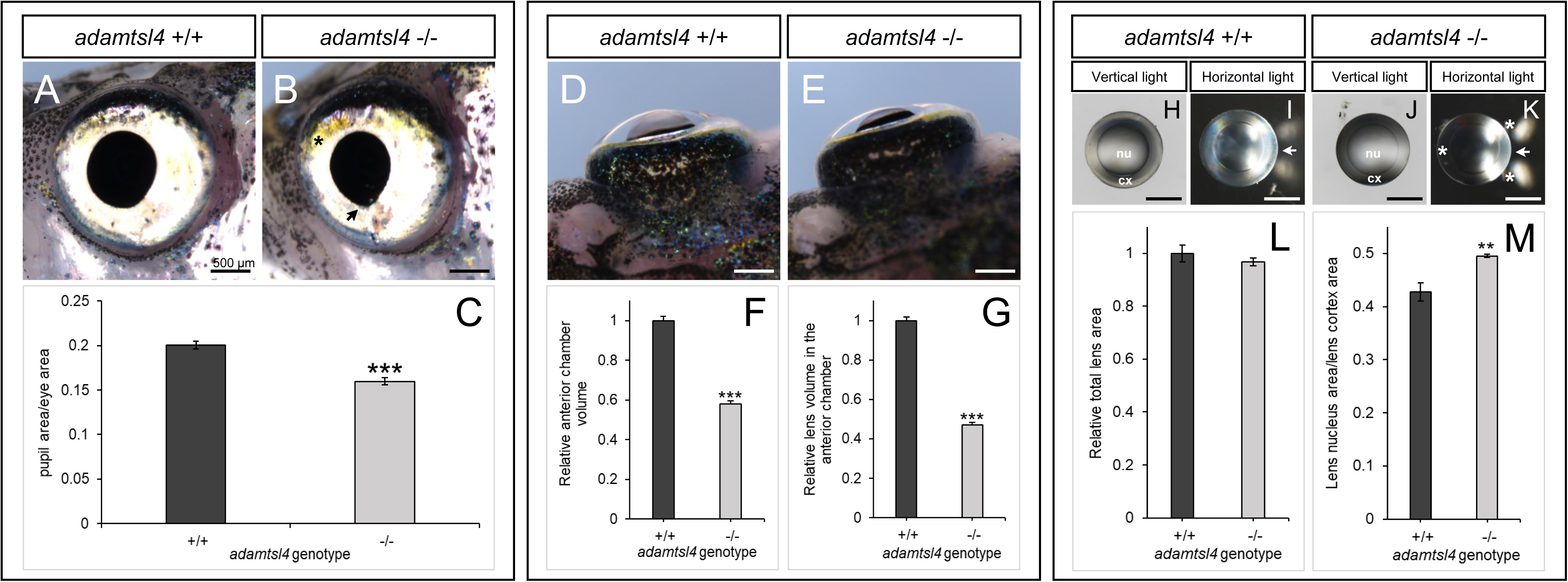
Morphological phenotype characterization of 5 months F3 *adamtsl4*-KO zebrafish. A-B. Lateral morphology of the eye. (C) Quantification of pupil area relative to the eye area. Sample size: wild-type, n=5; *adamtsl4*-KO, n=9. D-E. Dorsal morphology of the eyes. F. Quantification of anterior chamber volume. G. Quantification of the visible lens volume within the anterior chamber. Sample size in F-G: wild-type, n=20; *adamtsl4*-KO, n=72. H-K. Lens morphology and light transmission. L. Quantification of total lens area. M. Quantification of lens nucleus area relative to the cortex area. Sample sizes: wild-type, n=4; *adamtsl4*-KO, n=. **: p < 0.01; ***: p < 0.001. Black asterisk: increased xanthophores. Black arrow: irregular pupil morphology. White asterisk: impaired light transmission in the lens. White arrow: direction of light. Cx, cortex; nu, nucleus.

The structural and optical properties of isolated lenses were evaluated as detailed in the Materials and Methods section. Under vertical illumination, the wild type lens appeared transparent, allowing clear distinction between the nucleus and cortex (Fig. 6H). When illuminated horizontally, light passed through to the opposite side of the lens, creating two oblique reflection spots on either side of the incoming light (Fig. 6I). The lens of KO zebrafish also appeared transparent under vertical illumination and showed the separation between the nucleus and cortex (Fig. 6J). However, under horizontal illumination, the half of the KO lens facing the incoming light was illuminated. Additionally, the KO lens exhibited an increased intensity of reflected light compared to the wild type (Fig. 6K). No significant differences were observed in lens size (measured as total area) between wild-type and KO zebrafish (Fig. 6L). However, KO zebrafish exhibited a 1.16-fold increase in the ratio of lens nucleus area to cortex area compared to wild-type (Fig. 6M). These observations are consistent with an anomalous lens insertion or ectopia lentis and altered lens structure associated with *adamtsl4* LoF.

Histological analysis of adult zebrafish eyes, using hematoxilin-eosin stained sections corresponding to the central part of the eyeball revealed that in contrast to the wild type (Fig. 7A, C), the KO zebrafish exhibited an abnormal, continuous dorso-ventral extension of the annular ligament, which ran parallel to the inner surface of the cornea (Fig. 7B, D and F, asterisk). Furthermore, the corneal epithelium in the KO zebrafish was thicker, with a swollen superficial cell layer (Fig. 7E, F). The overall corneal thickness was increased by approximately 20% compared to the wild type animals (Fig. 7G). Additionally, an apparent increased separation between the lens fiber layers was observed in the KO zebrafish (Fig. 7D, arrows).

**Fig. 7.**
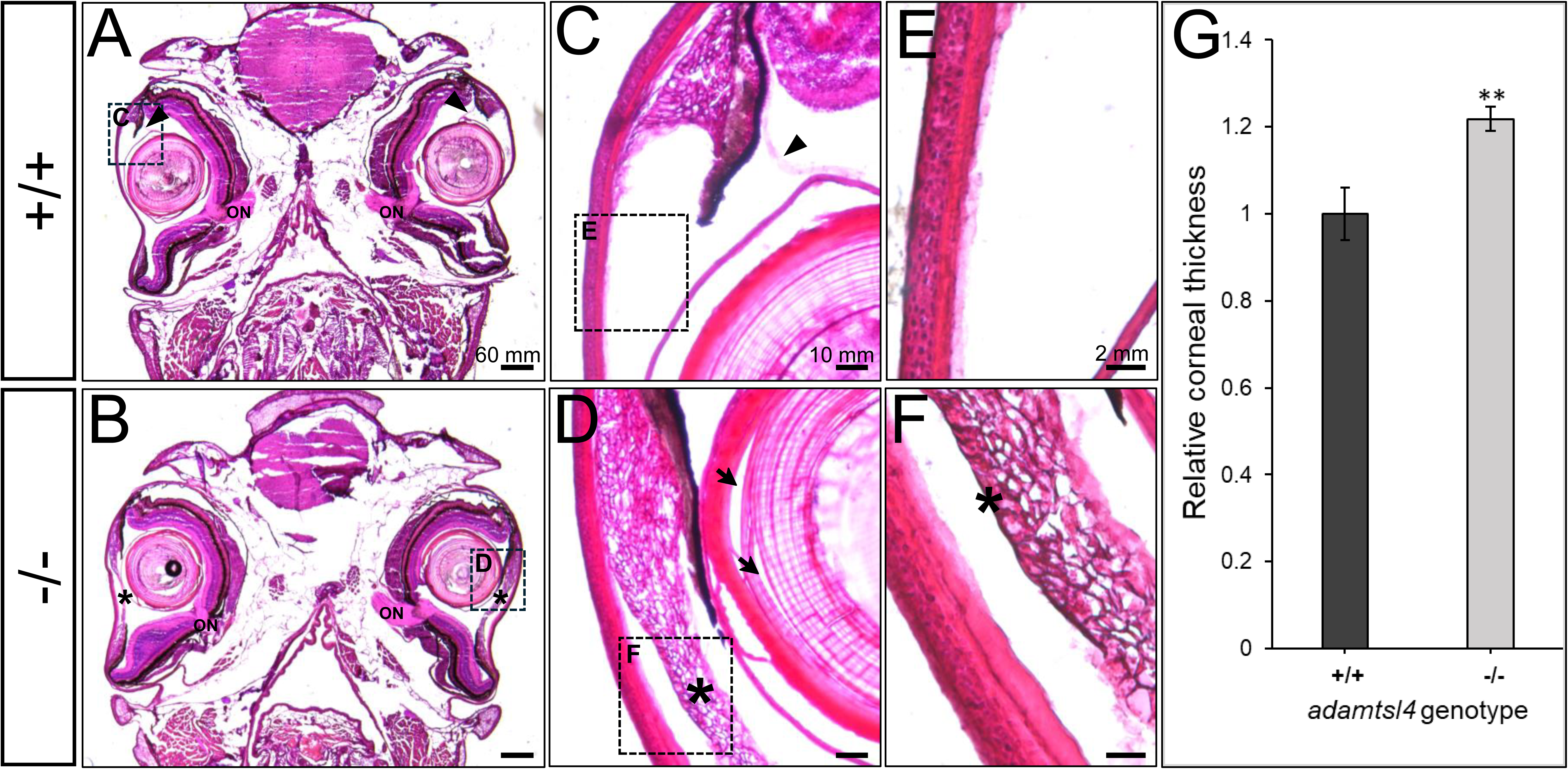
Histological analysis of hematoxylin-eosin-stained tissue sections from adult *adamtsl4* KO zebrafish (5 months). Transverse tissue sections were prepared as described in the Materials and Methods section. Representative photographs of wild type (+/+) and KO (-/-) zebrafish are shown. A-B. Tissue sections from the central part of the eyeball highlighting the presence of the optic nerve (ON). C-D. Anterior segment. E-F. Corneal structure. G. Quantitative analysis of corneal thickness. **: p<0.01. Rectangles indicate areas magnified in subsequent panels. Arrows: separation of lens fibers; Asterisks: abnormal annular ligament.

The ultrastructure of the eyes of adult wild type (Fig. 8A, C, E, G, I, K and M) and KO (Fig. 10B, D, F, H, J, L and N) zebrafish was examined using TEM. Like observations in KO larvae, the guanine crystals within the iridophores of adult KO eyes were enlarged and disordered compared to wild type eyes (Fig. 8A and B). Annular ligament cells exhibited increased electron density of glycoprotein-like deposits, as well as a greater number, size and electron density of intracellular inclusions (Fig. 8C and D; Gly and In, respectively). Corneal epithelial cells were swollen, with necrosis evident in the most superficial layer (Fig. 8E and F, asterisk), contributing to the observed increase in corneal thickness. In wild type zebrafish, the corneal stroma lamellae were tightly packed and highly ordered, with collagen fibrils in adjacent lamellae arranged orthogonally and presenting uniform size (Fig. 8G). Additionally, there is very little elastic tissue observed between the collagen fibers (Fig. 8G, arrows). In contrast, KO zebrafish displayed disorganized lamellae with irregular shape and size, along with an increased amount of elastic tissue interspersed between the collagen fibers (Fig 8H, arrows). The corneal endothelium was in contact with the abnormal annular ligament (Fig. 8H, arrowhead). Abnormal intercellular junctions, leading to large intercellular separations, were observed in the lens epithelium and lens fibers interphase (Fig. 8I-J, asterisks). The lens fibers also presented cellular junctions abnormalities, consisting of increased intercellular spaces (Fig. 8K-L, asterisks) surrounding gap junctions (Fig. 8K-L, white arrows). Normal autophagic vacuoles were present in the lens epithelium (Fig 8J, arrowhead). Abnormal intercellular separations were also identified at the interphase of the iris pigment epithelium and neuroretina (Fig. 8M and N, asterisk).

**Fig. 8.**
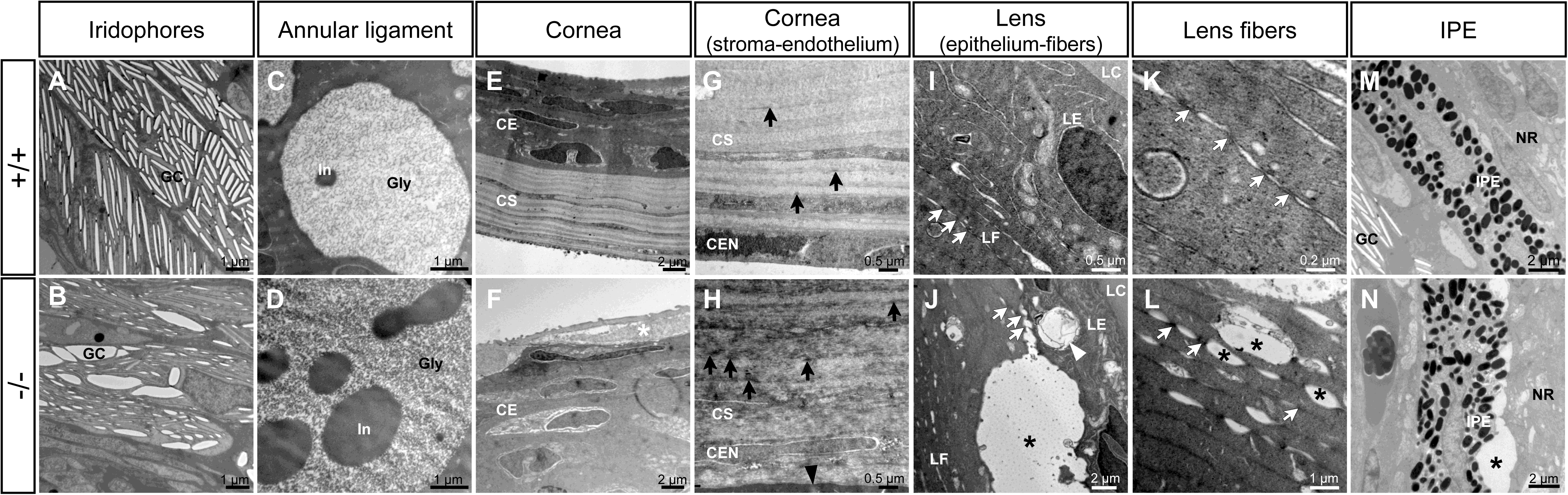
Transmission electron microscopy of the ocular anterior segment structures in adult (3-months-old) F3 *adamtsl4* KO zebrafish. Tissue sections were prepared as described in the Materials and Methods section. Representative images of wild type (+/+) and KO (-/-) zebrafish are shown. A-B. Guanine crystals (GC) in iridophores. C-D. Glycoprotein aggregates (Gly) and cytoplasmic inclusions (In) in annular ligament cells. E-F. Cornea showing the thickened corneal epithelium (CE) and the necrotic superficial epithelial cell (white asterisk) in the KO zebrafish. G-H. Corneal stroma (CS) and endothelium (CEN). Note the contact between the CEN with the abnormal annular ligament in the KO zebrafish (black arrowhead). I-J. Lens epithelium (LE)-fiber (LF) interphase. K-L. Lens fibers. Gap junctions (white arrows). M-N. Iris pigment epithelium (IPE)-neuroretina (NE) interphase. Black asterisks: abnormal intercellular separations.

### 3.4. Visual function assessment in adult KO zebrafish using the optokinetic response assay

The optokinetic response (OKR) assay was performed on 11-month-old *adamtsl4* KO F3 adult fish. We measured the angle of eye gaze over time in response to a rotating stimulus. In wild type zebrafish we observed a consistent pattern consisting of downward slow phases followed by upward rapid phases (Figure 9A). In contrast, KO zebrafish exhibited remarkable different responses to the same stimulus, characterized by small, irregular, or nearly absent eye movements (Fig. 9B). These findings indicate that the established KO line presents anomalies in visual function.

**Fig. 9.**
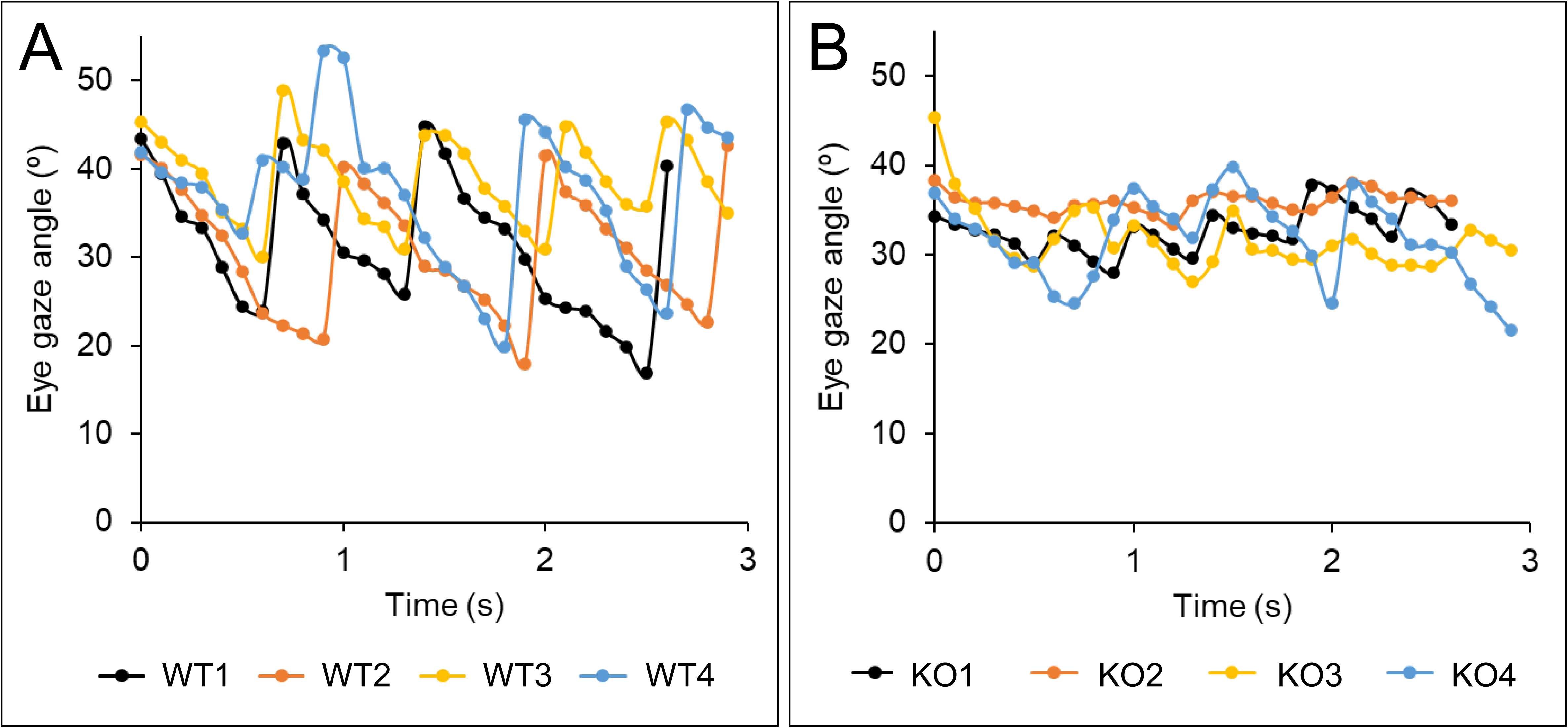
Optokinetic response in adult zebrafish at 8 months based on frame-by-frame eye angle measurements. A. Wild-type zebrafish. B. *adamtsl4*-KO zebrafish. Films were analyzed using Image-Pro Plus software to determine eye angles across individual frames. Data points represent eye angles measured in a representative analysis of four fish per group.

### 3.5. Differential gene expression analysis

Whole-transcriptome sequencing was carried out to evaluate differentially expressed genes (DEGs) associated with *adamtsl4* LoF in 6 dpf KO larvae (lethal and viable phenotypes) and in eyes from adult (5 months) KO zebrafish. Wild type specimens of the same age were used as controls. Two independent biological replicates of each experimental group were analyzed as indicated in the Materials and Methods section. Each mutant transcriptome was compared with the corresponding wild type transcriptome. After excluding unmapped genes and genes with more than one zero read count, 24,342 genes (out of 37,369 total genes, with 13,027 excluded) and 21,306 genes (out of 37,369 total genes, with 16,063 excluded) were retained for the statistical analysis of KO larvae and adult eye transcriptomes, respectively. All experimental replicates showed high homogeneity, as confirmed by the correlation matrix of samples calculated using Pearson’s coefficient (Supplementary Fig. S4A and C). We assessed the consistency of DEG patterns across replicates by visualizing hierarchical clustering in a heatmap. The results revealed similar DEG clusters within replicates of the same experimental group, indicating that the identified gene expression patterns are highly reproducible and consistent across experiments (Supplementary Fig. S4A and D).

By applying a differential expression filter (log2 fold change ≥ |1|, p < 0.05), we identified 1,028 DEGs in 6 dpf lethal KO larvae compared to wild-type larvae, with 879 upregulated and 149 downregulated genes. In 6 dpf viable wild type-like KO larvae vs. wild-type larvae, we found 133 DEGs, including 96 upregulated and 37 downregulated genes. For KO adult eyes vs. wild-type eyes, we observed 3,388 DEGs, with 1,255 overexpressed and 2,133 underexpressed genes. For the initial functional evaluation of DEGs, we selected the top 50 DEGs from each comparison, comprising the 25 most upregulated and the 25 most downregulated genes. Among down-regulated genes in lethal KO larvae, the majority (13) were of uncharacterized function, and the rest were involved in ion transport and channel activity, metabolism and transport, immune response and lens and eye development (three genes in each group, Supplementary Table S2 and Fig 10A). Notably, 11 overexpressed genes were involved in ECM and cell adhesion, six in the inflammatory response and immune function, three in stress response and protein folding, two in developmental signaling and transcription regulation. Only one gene was of uncharacterized function another was associated with cytoskeletal organization, and one was involved in metabolism (Supplementary Table S2 and Fig. 10B).

**Fig. 10.**
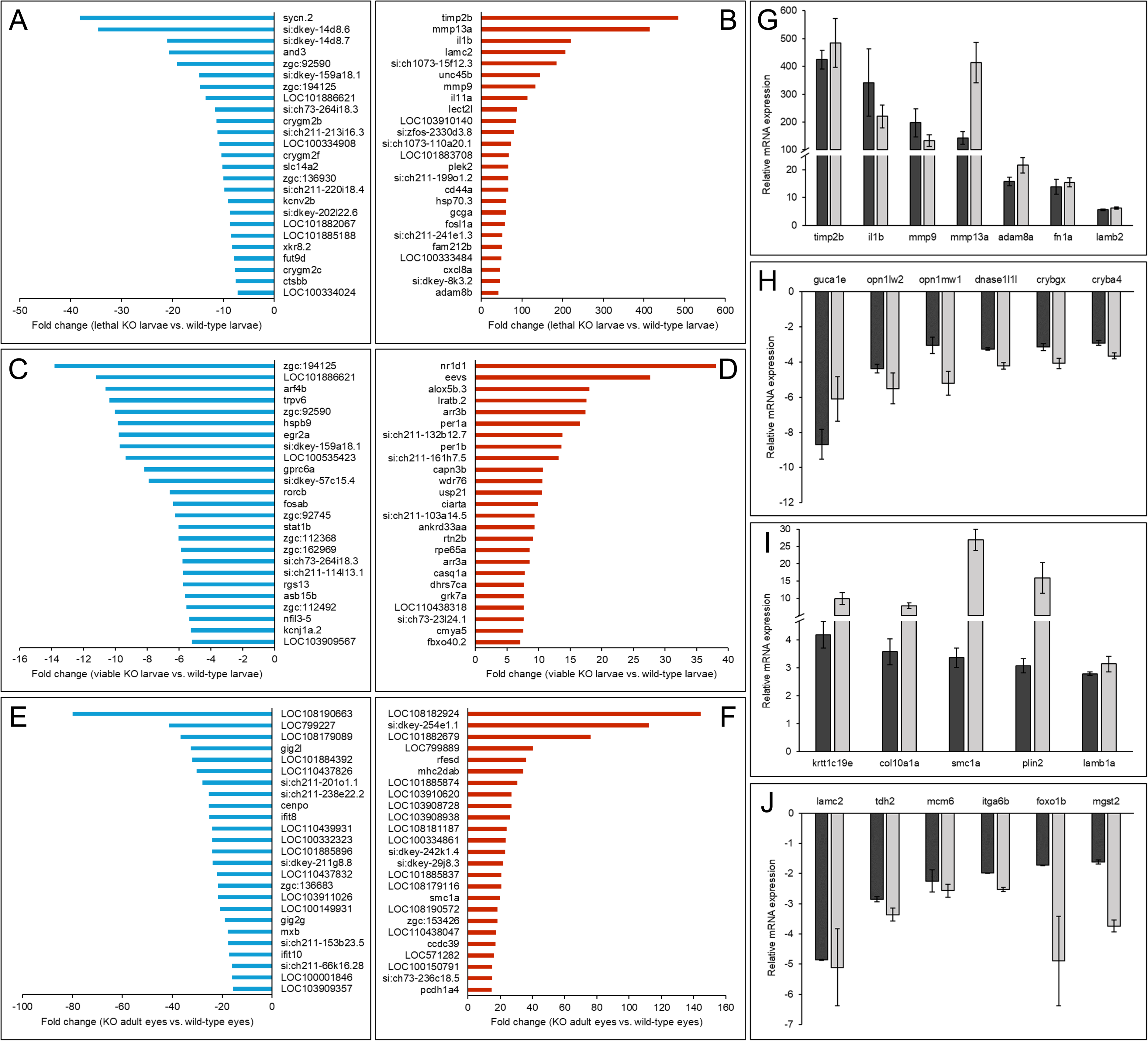
Top 50 DEGs in adamtsl4 KO zebrafish identified by RNA-seq analysis and confirmation by RT-qPCR of the differential expression of selected genes. A-B. Downregulated genes and upregulated genes, respectively, in 6 dpf lethal *adamtsl4* KO larvae vs. wild type larvae. C-D. Downregulated genes and upregulated genes, respectively, in 6 dpf wild type-like *adamtsl4* KO larvae vs. wild type larvae. E-F. Downregulated genes and upregulated genes, respectively, in eyes of 5-month *adamtsl4* KO zebrafish vs. wild type zebrafish. G-H. Confirmation by qRT-PCR of differential gene expression of selected upregulated and downregulated genes, respectively, in lethal *adamtsl4* KO larvae vs. wild type larvae. I-J. Confirmation by qRT-PCR of differential gene expression of selected upregulated and downregulated genes, respectively, in eyes of 5-monts *adamtsl4* KO zebrafish vs. wild type zebrafish. Aliquots of RNA preparations used for transcriptomic analyses were used as templates in qRT-PCR that was carried out in triplicate.

In viable wild type-like KO larvae we observed overexpression of a different set of genes, with roles in vision and phototransduction (5 genes), muscle-related functions (5 genes), circadian rhythm regulation (4 genes), lipid and retinoid metabolism (3 genes) and protein modification and degradation (5 genes). Three genes were of uncharacterized function (Fig. 10C, Supplementary Table S3). Downregulated genes participate in transcription (5 genes), cellular signaling (4 genes), cellular transport (2 genes) and protein modulation/degradation (2 genes). Most genes had uncharacterized functions (12 genes) (Fig. 10D, Supplementary Table S3).

Finally, in eyes of adult KO zebrafish most of the top 50 DEGs were of unknown function, but we detected upregulated expression of genes that participated in cell adhesion (*pcdh1a4*), cilia structure and function (*ccdc39 and cfap70*), chromosomal cohesion (*smc1a*) and immunity (*mhc2dab*), while five interferon response-related genes were underexpressed (Fig. 10E-F, Supplementary Table S4).

### 3.6. Functional Enrichment of DEGs

All DEGs genes identified in each experimental group were subjected to functional pathway analysis using the bioinformatic tool ShinyGO, selecting the the 20 most significant biological, grouped into functional classes and ranked by FDR. Consistent with the initial results, analysis of the 20 most significantly enriched biological processes in 6 dpf lethal KO larvae revealed that the majority were related with two main functional activities: (1) immune and stress-related responses and (2) development processes of the eye and other organs (Supplementary Table S5).

In contrast, the same ShinyGO analysis of the transcriptome of viable wild type-like larvae showed that the 20 most enriched biological processed were related to circadian rhythm and rhythmic processes, guanylate cyclase activity, DNA replication and repair and mesenchyme migration and morphogenesis (Supplementary Table S6).

In adult KO zebrafish eyes, the DEGs were associated with biological processes such as ECM (genes of the transglutaminase family that form covalent bonds between extracellular proteins), cell adhesion, development and morphogenesis and metabolism (Supplementary Table S7). These findings suggest that *adamtsl4* LoF, depending on the genetic background, can result in larval lethality and altered expression of genes involved in organ development, immune responses, and stress pathways. In the eyes of adult zebrafish with genetic backgrounds that support embryonic viability, *adamtsl4* disruption impacts circadian rhythms, visual mechanisms, DNA-related processes, and metabolic pathways. These effects may contribute to the observed ocular phenotypes.

### 3.7. Validation of the Transcriptomic Analysis

To validate the transcriptomic results, RT-qPCR was performed on 13 genes representing the main DEG functional classes identified in lethal KO zebrafish larvae. These included genes related to ECM and cell adhesion (*timp2b*, *mmp9*, *mmp13a*, *adam8a*, *fn1a*, and *lamb2*), lens structure (*crybgx* and *cryba4*), inflammation and immune response (*il1b*), visual function (*guca1e*, *opn1lw2*, and *opn1mw1*), and DNA processes (*dnase1l1l*). The RT-qPCR results corroborated the RNA-seq findings, revealing significant overexpression of the ECM and cell adhesion-related genes, with fold changes ranging from approximately 10-fold (*lamb2*) to more than 400-fold (*timp2b*) (Fig. 10G). In contrast, the genes associated with the other three functional classes were significantly underexpressed in the KO larvae, showing negative fold changes ranging from -4 to -8 (Fig. 10H).

RT-qPCR analysis of 11 selected DEGs from the eyes of adult KO zebrafish also validated the transcriptome results. However, the gene expression differences observed by PCR were higher than those detected by RNA-seq (Fig. 10I). This discrepancy is likely due to methodological differences between the two techniques. RT-qPCR is widely considered the gold standard for quantifying gene expression differences, offering higher sensitivity and specificity than RNA-seq. The selected genes represent the main functional groups: ECM and cell adhesion (*krtt1c19e*, *col10a1a*, *lamb1a*, *lamc2* and *itga6b*), metabolic processes (*plin2*, *tdh2* and *mgst2*) and regulation of DNA replication (*smc1a* and *mcm6*) and gene expression (*foxo1b*). Genes related to ECM and cell adhesion showed a significant upregulation, with expression increasing approximately 5-to 25-fold in the eyes of adult KO zebrafish. In contrast, genes from the other categories were significantly downregulated (Fig. 10J).

## 4. Discussion

*ADAMTSL4* LoF *c*ause a spectrum of ocular and non-ocular conditions, such as ectopia lentis [3, 4], iris abnormalities and craniosynostosis [1]. Despite extensive investigations, the precise biological function of this gene and the molecular mechanisms driving these phenotypes remain largely unknown. Developing animal models could significantly enhance our understanding of the physiological roles and pathogenic pathways associated with ADAMTSL4. To the best of our knowledge, we report the first KO zebrafish line generated to study the molecular base of *adamtsl4-*related diseases. The developed zebrafish line harbored a deletion in the *adamtsl4* gene that was predicted to result in a frameshift and in the presence of a premature stop codon in the mutant protein (p.(Gln78Hisfs*127)), justifying a complete LoF. In fact, we demonstrated by RT-qPCR a significant reduction in *adamtsl4* mRNA levels, which correlated with the number of mutant alleles. This reduction is likely the result of the NMD mechanism, triggered by the presence of premature termination codons [24]. Furthermore, the translation of the residual 25% mRNA observed in KO zebrafish would produce a truncated protein missing more than 90% of its amino acid sequence, making it very likely to be completely inactive.

We observed that approximately 60% of F3 KO larvae manifested no severe abnormalities, while the remaining 40% presented lethal malformations, demonstrating the existence of incomplete penetrance. This finding highlights the involvement of *adamtsl4* in normal zebrafish embryonic development. Moreover, it suggests that the functional disruption of this gene can lead to lethality depending on the genetic background. Although lethality due to ADAMTSL4 loss-of-function (LoF) mutations has not, to our knowledge, been reported in humans, our findings suggest that such mutations could be incompatible with survival in fetuses with a genetically susceptible background. This might lead to spontaneous abortions that have not been attributed to disruptions in this gene. The reported phenotypic variability among patients with *ADAMTSL4* mutations [6] also supports that the phenotypic outcome of *ADAMTSL4* LoF may be influenced by modifier genetic and/or environmental factors. Ocular histology revealed clear ECM alterations of anterior segment structures (iris, cornea and lens) that were more severe in the eyes of the lethal larvae. The main ECM alterations, e.g., thickening of both lens capsule and corneal stroma (the latter being accompanied with abnormal deposition of collagen fibers) were also present in the eyes of adult KO zebrafish. The zonular fibers were similar in the eyes from wild type and KO zebrafish, at least at the optical microcopy level. Interestingly, the presence of intercellular separations, observed between corneal epithelial cells and between the lens epithelium and peripheral lens fibers as well as between the RPE and the neuroretina, indicate that *adamtsl4* LoF also alters cell junctions, implying a crucial role for the adamtsl4 protein in maintaining cell-cell adhesion. These RPE alterations may explain the retinal detachment described in some patients with ADAMTSL4 mutations [4]. Notably, gap junctions appeared severely compromised, particularly in the lens of KO zebrafish. Supporting this hypothesis, the transcriptomic analysis of the KO specimens revealed dysregulation of several genes involved in cell adhesion, as discussed in subsequent paragraphs.

The macroscopic ocular phenotypes observed in the adult *adamtsl4* KO zebrafish included a decreased relative area of both the pupil and lens in the anterior chamber, altered pupil morphology, corneal thickening and abnormal cranial morphology with irregular cranial sutures. These findings indicate that the zebrafish model replicates key human phenotypes associated with *ADAMTSL4* LoF, such as ectopia lentis et pupillae, increased corneal thickness and craniosynostosis [1, 2]. The underlying cause of these phenotypes is the disruption of the biological function of *ADAMTSL4,* which is not yet fully understood. This protein is known to regulate the assembly and stability of fibrillin-containing microfibrils [11]. It also plays a role in elastic fiber development by serving as a scaffold for tropoelastin deposition and modulating microfibril interactions with other ECM proteins [25]. In the ocular ciliary zonule ADAMTSL4 binds fibrillin-1 microfibrils and enhances their deposition within the ECM [10]. Microfibrils are important for tissue homeostasis as they act as reservoirs for latent TGF-β and also interact with bone morphogenetic proteins and with cell surface receptors such as the integrins [26, 27] and syndecans [28]. Therefore, dysregulation of ADAMTSL4 can interfere with the assembly or maintenance of the zonular microfibrils, leading to a weakened zonule and lens dislocation, as well as to abnormal organization of elastic fibers in the corneal stroma of KO zebrafish. Additionally, ADAMTSL4 is involved in maintaining the structure and stability of cranial sutures [2]. *Adamtsl4* LoF may alter TGF-β bioavailability and ECM remodeling, which contributes not only to ocular and cranial phenotypes but also to the broader developmental abnormalities observed in this study. Supporting this idea, *ADAMTSL4* variants have been found to phenocopy TGFβ receptor mutations in craniosynostosis patients, suggesting that *ADAMTSL4* LoF disrupts TGFβ signaling required for regulating cranial suture patency. This provides an hypothetical mechanism for the abnormal bone fusion characteristic of craniosynostosis [2].

The lens of adult KO zebrafish exhibited altered optical properties, likely resulting from observed disruptions in the ECM, cell junctions, and crystallin expression. Lens transparency and normal refractive index depend on precise lens fiber alignment, minimal intercellular space, and high concentrations of cytoplasmic crystallins, among other factors. Any disruption of the fine organization of the lens fibers and/or modifications of their proteins can compromise transparency [29]. In our study, we detected increased intercellular spaces surrounding gap junctions and differential expression of several crystallin genes. In adult KO zebrafish, genes such as *crygm4*, *hspb3* (*heat shock protein beta-3, alpha-crystallin-related*), and hspb11 *(heat shock protein beta-11, alpha-crystallin-related*) were differentially expressed. Additionally, lethal KO larvae showed changes in *crygm2c*, *crygm2f*, and *crygm2b* expression. These findings not only explain the altered optical properties observed in adult KO zebrafish lenses but also suggest potential structural changes in the lenses of patients with *ADAMTSL4* mutations that may explain the cataracts present in some of these patients [4].

Some characteristics of the observed ocular phenotypes align with those reported in mice with disrupted *Adamtsl4,* obtained through an ethylnitrosourea mutagenesis screen. These mutant mice presented detachment of zonular fibers from the lens [15]. Interestingly, the mice also presented defects in the RPE, marked by age-dependent dedifferentiation. Notably, consistent with our findings, these mice also displayed downregulation of *Rpe65,* an RPE-specific gene critical for photoreceptor function and visual perception. However, the study did not report any data on lethality or non-ocular abnormalities.

In the human eye, *ADAMTSL4* is expressed in the stroma of the iris and ciliary body, as well as in the ciliary processes and RPE [9, 11, 30]. This protein is also present in the lens epithelium and fibers, corneal epithelium and endothelium, as well as the iris stroma, iris fibroblasts and iris sphincter muscle [9]. In zebrafish larvae (6 dpf), *adamtsl4* is detected in the cornea, lens epithelium and periocular mesenchyme, which contribute to the morphogenesis of the iris and anterior chamber angle structures [9]. The expression of adamtsl4 in altered ocular anterior segment structures observed in KO animals and patients with mutations in this gene underscores its involvement in the observed phenotypes.

Although *ADAMTSL4* is widely expressed thorughout the body (<u>GTEx Portal</u>, accessed on 12/01/2024), no phenotypes in other tissues or organs have been linked to its functional alteration. This suggests compensatory molecular or proteomic mechanisms in non-ocular tissues mitigate the impact of ADAMTSL4 dysfunction, as previously proposed [2]. Consistent with this hypothesis, it can be speculated that mechanisms involved in this genetic or molecular compensation may also contribute to the phenotypic variability observed in ADAMTSL4-related diseases.

Our transcriptomic analyses have provided new insights into molecular mechanisms underlying the phenotypes associated with *adamtsl4* LoF mutations, as well as into the biology of this gene. One of the most interesting findings revealed by functional analysis of DEGs was the significant overexpression of genes related to ECM components and cell adhesion. Additionally, we observed dysregulation in genes associated with immune and stress responses, as well as organ development, in lethal knockout (KO) zebrafish larvae, also confirmed by RT-qPCR. While establishing direct cause-and-effect relationships among DEGs is challenging, we hypothesize that the disruption of ECM maintenance and cell adhesion gene expression during zebrafish development is likely the primary effect of *adamtsl4* inactivation. In contrast, the altered expression of most other genes may reflect secondary responses to the primary defects in embryonic ECM structure. Supporting this idea, the transcriptome of wild type-like KO larvae showed no significant changes in the expression of immune or stress-related response genes. This finding also suggests that the secondary responses associated with *adamtsl4* LoF, are dependent on the genetic background. Validation by RT-qPCR confirmed a significant upregulation of selected ECM and cellular adhesion-related genes, with expression levels in some cases exceeding those of the control group by one or even two orders of magnitude. The selected genes encoded a matrix metalloproteinase inhibitor (*timp2b*), three metalloproteinases (*mmp9*, *mmp13a*, and *adam8a*), and two cell adhesion-related proteins (*fn1a* and *lamb2*). The collective dysregulation of these genes also supports the hypothesis that *adamtsl4* inactivation likely disrupts the delicate balance of ECM remodelling, cell adhesion, and signalling during embryogenesis. Such disruptions may lead to severe developmental abnormalities and larvae lethality. These findings also underscore the key role of *adamtsl4* in regulating ECM dynamics and cell adhesion processes. In accordance with this idea, two ADAMTS family members, ADAMTS6 and ADAMTS10, regulate focal adhesions, epithelial cell-cell junction formation, and microfibril deposition [31].

Interestingly, transcriptome analysis of wild type-like KO larvae revealed specific DEGs. These included genes involved in circadian rhythm and rhythmic processes, guanylate cyclase activity, DNA replication and repair, as well as mesenchyme migration and morphogenesis. These changes, absent in lethal KO larvae suggests that these DEGs might contribute to larval survival by partially compensating for *adamtsl4* LoF and/or driving specific phenotypic alterations. The differential expression of circadian rhythm-related genes (*nr1d1*, *per1a*, *per1b* and *ciarta*) in wild type-like KO zebrafish may influence various physiological processes, including metabolism, inflammation, immunity, and cardiovascular functions [32–35]. These changes could play a role in improving survival outcomes.

The transcriptome analysis of adult KO zebrafish eyes also revealed significant changes in the expression of genes involved in ECM and cell adhesion, as well as in metabolism, development and morphogenesis. Regarding the first group of DEGs, several were involved in cell junctions, potentially explaining the abnormal intercellular separations observed in the cornea, lens and RPE of adult KO zebrafish. Notably, among the identified genes, several encode claudins (*cldnb*, *cldn5a*, *cldn5b*, *cldne*, *cldn19*, and *cldna*), which are key proteins in tight junction formation. Tight junctions create barriers in epithelial and endothelial layers, regulate paracellular transport, and maintain epithelial cell polarity [36]. Tight junctions, along with adherens junctions and desmosomes, form multi-protein complexes that mediate the adhesion of corneal epithelial cells at their apicolateral surfaces [37]. Additionally, three genes (*cdh27*, *cadm3*, and *ncam3*) encode proteins involved or predicted to be involved in cell adhesion (Zebrafish Information Network (ZFIN), University of Oregon, Eugene, OR 97403-5274; URL: http://zfin.org/; [accessed on 12/02/2024]). The group of differentially expressed genes (DEGs) also includes two plakophilin genes (*pkp2* and *pkp3a*) and one desmocollin gene (*dsc2l*), essential for the stability of desmosomal junctions [38, 39](Zebrafish Information Network (ZFIN), University of Oregon, Eugene, OR 97403-5274; URL: http://zfin.org/; [accessed on 12/02/2024]). Interestingly, the expression of *jupa* (*junction plakoglobin a*) and *igsf5a* (*Immunoglobulin Superfamily Member 5a*) was also altered. The human ortholog of the first gene is involved both in desmosomes and adherens junctions [40], while the second participates in bicellular tight junction and cell surface adhesion [41]. Furthermore, several hemidesmosome integrins (*itga2b*, *itgb8*, *itga1*, *itgam*, and *itgae.1*), a focal adhesion kinase (*ptk2ba*), laminins (l*amb2l* and *lamb1a*), and thrombospondins (*thbs4b*), along with associated proteins (*ntn4*, *mag*, *spp1*, *postnb*, *frem1a*, and *frem2a*), were found to be dysregulated. These components are crucial for interactions between the basement membrane and the extracellular matrix (ECM). Different protocadherins (*pcdh2aa15*, *pcdh2ab9*, *pcdh2ac*, *pcdh1g3*, *pcdh12*, and *pcdh1a4*), which play roles in neural development and non-classical adhesion [42], were also altered. Additionally, junction-cytoskeleton linking proteins (*macf1b*, *ctnnal1*, and *lurap1)* [43, 44] and other cell adhesion molecules (*vcam1b* and *cd44a*) showed changes in expression. These proteins participate in processes like immune cell interactions [45] and in maintaining organ and tissue structure through cell-cell and cell-matrix interactions [46]. Collectively, these findings suggest that loss of function (LoF) mutations in *adamtsl4* may disrupt cell-cell and cell-ECM adhesion, as well as impair ECM-related signal transduction processes, contributing to the observed ocular phenotypes, particularly the distinctive intercellular separations observed in different parts of the eye of KO zebrafish.

The differential expression of genes in the eyes of adult KO zebrafish, representing the main functional groups, was also confirmed by RT-qPCR. Particularly interesting, the five genes selected ECM and cell adhesion-related genes were significantly overexpressed, demonstrating that adamtsl4 LoF impacts cell adhesion processes. Among these genes, *krtt1c19e* encodes a keratin expressed in basal epidermal keratocytes that likely contributes to the cytoskeletal framework of epithelial cells [47], as well as to epithelial integrity trough the attachment of keratin intermediate filaments to desmosomes [48]. Collagen (*col10a1a*) and two laminins (*lamb1a* and *lamc2)* are key components of the ECM and basement membrane [49], which are essential for cell adhesion [50] The final selected gene, *itga6b*, encodes an integrin that is part of the laminin receptor complex [51].

In conclusion, our study reinforces the role of adamtsl4 LoF in ectopia lentis *et pupilae*, cataracts, retinal detachment and craniosynostosis. Moreover, it provides new insights into the gene’s biological functions in embryonic development and the regulation of cell junctions. Additionally, this research establishes a valuable animal model for investigating *ADAMTSL4*-related diseases and their poorly understood underlying mechanisms.

## Acknowledgements

We thank María-José Cabañero for her exceptional technical assistance and the Centro Nacional de Microscopía Electrónica, especially Dr. María-Luisa García-Gil for her excellent help with electron microscopy. This study was funded by grants from the “Instituto de Salud Carlos III” (Award Numbers: RD16/0008/0019 and PI19/00208. Recipient: Julio Escribano); the “Ministerio de Ciencia, Innovación y Universidades (MICIU)/Agencia Estatal de Investigación (AEI) y FEDER, UE” (Award Number: PID2023-149543OB-C22. Recipients: Jose-Daniel Aroca-Aguilar and Julio Escribano); the “Consejería de Ciencia y Tecnología de la Junta de Comunidades de Castilla-La Mancha” (Award Numbers: SBPLY/17/180501/000404 for Julio Escribano and SBPLY/23/180225/000206 for José-Daniel Aroca-Aguilar and Julio Escribano); and the “Universidad de Castilla-La Mancha” (Award Numbers: 2022-GRIN-34136. Recipient: Julio Escribano; and 2020-PREDUCLM-16605. Recipient: Angel Tevar).

