## Supplementary information for "Zebrafish *adamtsl4* knockout recapitulates key features of human *ADAMTSL4*-related diseases: a gene involved in extracellular matrix organization, cell junctions and development"

[1]Laboratory of Human Molecular Genetics, Albacete Medical School, Castilla-La Mancha University, Albacete, Spain
[2]Institute of Biomedicina, Castilla-La Mancha University (IB-UCLM) Albacete, Spain
[3]Ramon Castroviejo Institute for Ophthalmic Research, Complutense University of Madrid, Madrid, Spain
[4]Health Research Institute of the Hospital Clínico San Carlos (IdISSC), 28040 Madrid, Spain
[5]Department of Immunology, Ophthalmology and ENT, Faculty of Optics and Optometry, Complutense University of Madrid, Madrid, Spain
[6]Current address: University Institute of Biotechnology and Biomedicine (BIOTECMED), Valencia University, Valencia, Spain
*Co-corresponding authors at*:* Laboratory of Human Molecular Genetics, Albacete Medical School, Castilla-La Mancha University, Albacete, Spain

**This file includes:**

- **Figures S1-S4**
- **Tables S1-S7**

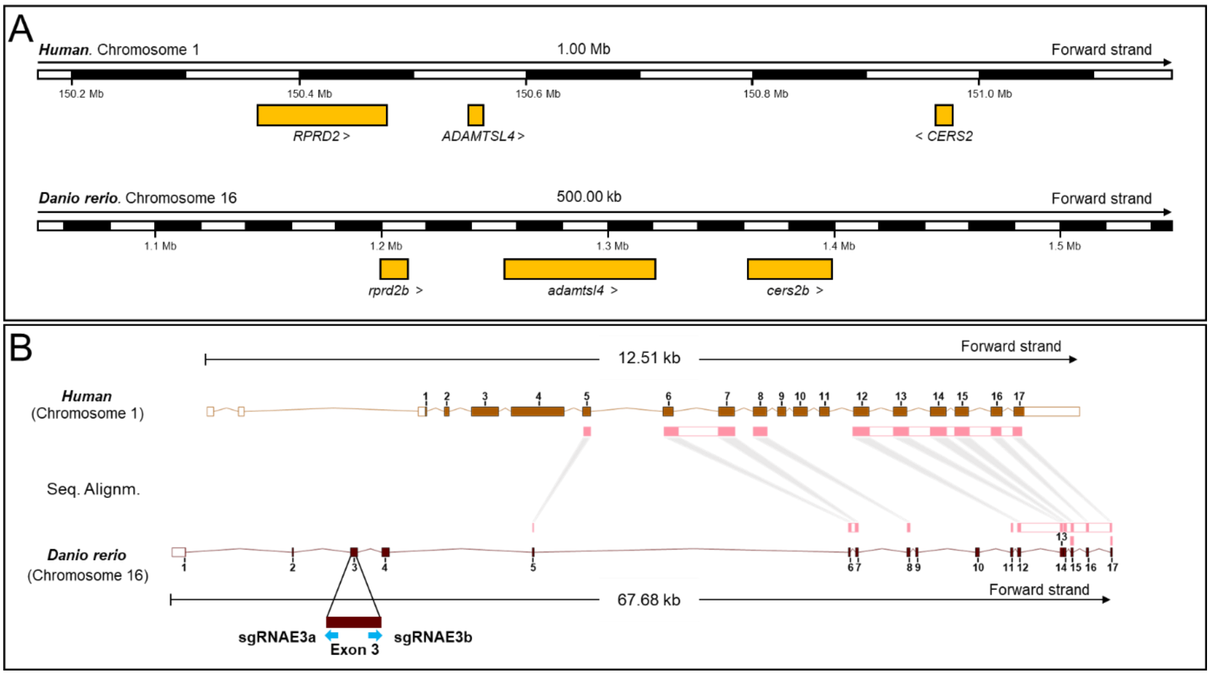


**Supplementary Fig. S1. Synteny and genomic alignment of human and zebrafish *ADAMTSL4* genes.**

A. Synteny analysis comparing the genomic regions surrounding the *ADAMTSL4* gene in humans and zebrafish. B. Genomic alignment of the human (ENSG00000143382) and zebrafish (ENSDARG00000063407) *ADAMTSL4* genes. The alignment was generated using the Ensembl Region Comparison Tool. Pink rectangles indicate conserved exons. The position of the two sgRNAs used to generate the adamtsl4 KO zebrafish are indicated by blue arrows.

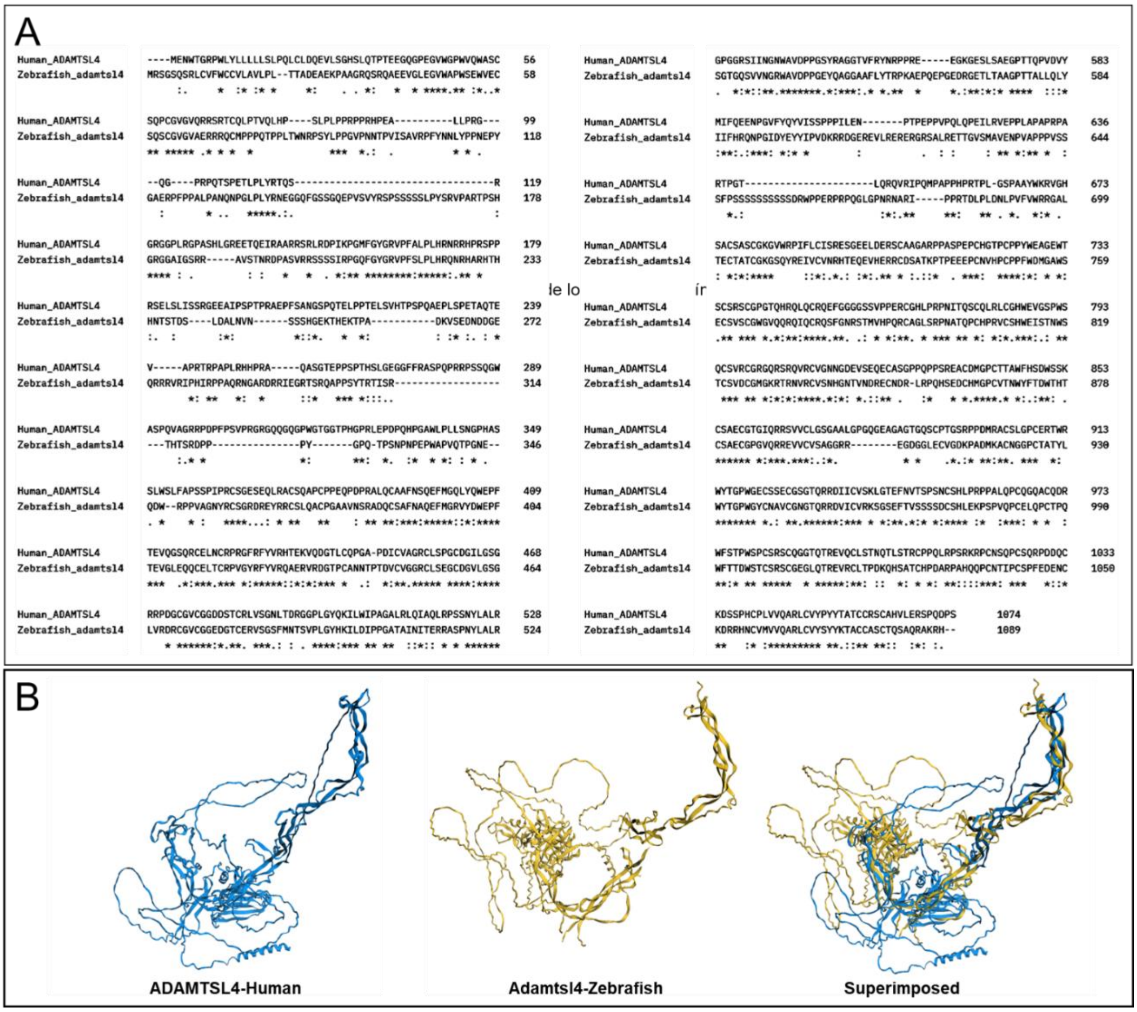


**Supplementary Fig. S2. Comparative analysis of human and zebrafish ADAMTSL4 proteins.**

A. Amino acid sequence alignment of human (ENSP00000358035) and zebrafish (ENSDARP00000087059) ADAMTSL4 proteins. Alignment was performed using Clustal Omega (https://www.ebi.ac.uk/jdispatcher/msa/clustalo). Asterisks (*) indicate positions with identical amino acids, two vertical dots (:) denote amino acids with similar chemical properties, and a single dot (.) represents positions with weak chemical similarity. B. Structural modeling of orthologous ADAMTSL4 proteins in humans (blue) and zebrafish (yellow) using deep learning and artificial intelligence methods. Protein structures were retrieved from the AlphaFold Protein Structure Database (https://alphafold.ebi.ac.uk/).

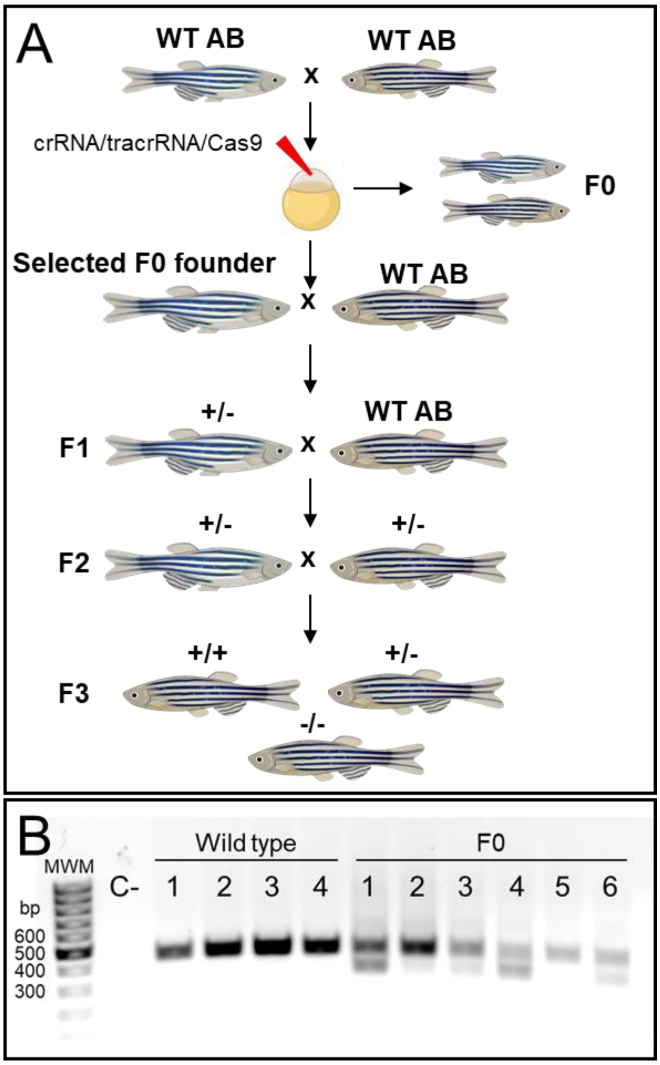


**Supplementary Fig. S3. Generation of the *adamtsl4*-KO zebrafish line using CRISPR/Cas9 genome editing.**

A. Schematic representation of the stepwise procedure followed to establish the KO line. F0 animals were raised to adulthood and crossed with wild-type AB zebrafish. Offspring were genotyped via PCR to confirm germline transmission of *adamtsl4* deletions (F0 founders). F0 founders were crossed with wild-type AB zebrafish to produce F1 heterozygotes. These heterozygotes were outbred to segregate potential off-target mutations, generating the F2 generation. F2 heterozygotes were inbred to produce the F3 generation. This schematic was created using the Biorender tool. B. Identification of *adamtsl4* mutant F0 zebrafish by PCR and agarose electrophoresis (1%). Representative samples include four wild types and six F0 animals. High and low molecular weight bands correspond to wild-type and mutant alleles, respectively.

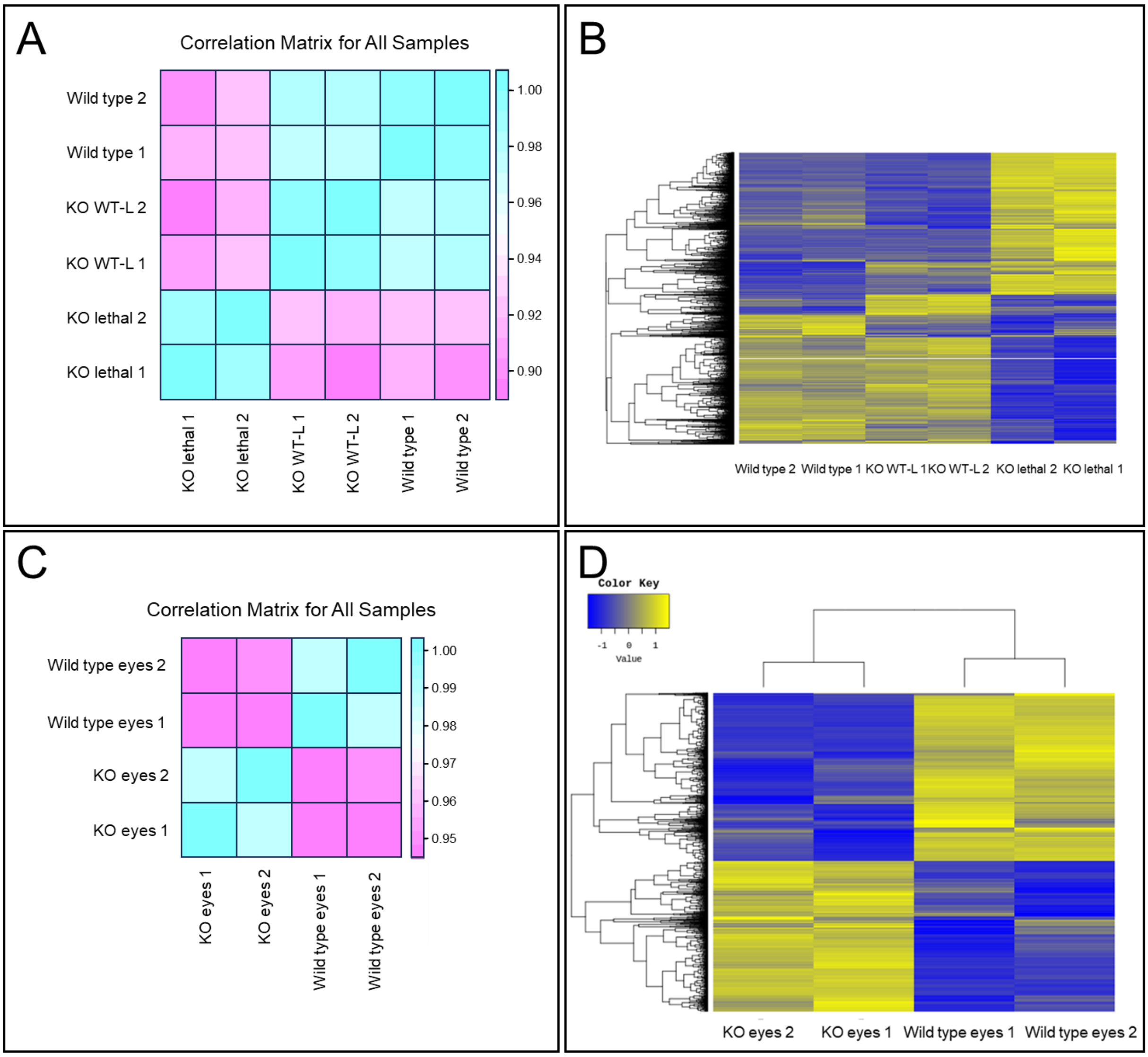


**Supplementary Fig. S4. Analyses of DEGs identified by RNA-Seq.**

A-B. DEGs identified in wil type-like adamtsl4 zebrafish larvae (6 dpf) compared to wild type larvae (6 dpf). C-D. DEGs identified in the eyes of adult adamtsl4 KO zebrafish compared to the eyes of wild type zebrafish of the same age. A and C. Correlation matrix of all samples, illustrating sample similarity based on Pearson's correlation coefficient of normalized expression values. The coefficient range ($-1 \leq r \leq 1$) indicates the degree of similarity, with values closer to 1 representing higher similarity. B and D. Heatmaps representing hierarchical clustering (Euclidean distance, complete linkage). Clustering shows relationships among genes and samples based on expression levels. The color key indicates normalized expression intensity: blue represents the highest expression levels, and yellow indicates the lowest.

**Supplementary Table S1.**
Primers sequences used in RT-qPCR.

| Gene | Primer | Sequence (5’ to 3’) |
|---|---|---|
| *adam8a* | Fw | CCTGCCTCCTTCAAGACCATTACC |
| | Rv | TGAGGCCCGCTGGGTTTC |
| *adamtsl4* | Fw | CATCCAGACGCCAGACCTGCAC |
| | Rv | CAGAGCCGAGCCTGAACCACC |
| *cryba4* | Fw | CTCCTTTAGACCAATCTCCTGCGCC |
| | Rv | CAGCTCGCCTTTACGGCCC |
| *crybgx* | Fw | CGACTGGACCTGACCTCTGA |
| | Rv | CCCCTGGAAGTTGTCATGTT |
| *col10a1a* | Fw | TCAATGAGCAGGACACAGTTTA |
| | Rv | GGCAATCAGGAATCCAGAGAA |
| *dnase1l1l* | Fw | CAGAGACGAACAAGACACTACAGTGCG |
| | Rv | CGGTTGAGCGGATTCCGGAACTATC |
| *ef1α* | Fw | CTGGAGGCCAGCTCAAACAT |
| | Rv | ATCAAGAAGAGTAGTACCGCTAGCATTAC |
| *fn1a* | Fw | TGGTGCTGAAGTTGATGCAGACCTC |
| | Rv | GTCTCAGGCACTCAATGGGACACTG |
| *foxo1b* | Fw | CCTGAACGGATGTGCAATGT |
| | Rv | CCACCTCTTCCCAAATGAGTATG |
| *guca1e* | Fw | GGACGAACTGCTGCTCATCTTCA |
| | Rv | CCAAGTCCTCTGCAGAGATCTCC |
| *il1b* | Fw | CAGTGCTCCGGCTACTTCTGC |
| | Rv | CCAGCTTTAATGGTCTCCAGAGACCC |
| *itga6b* | Fw | CTGTATTTCCAGAACGCAAAGTAG |
| | Rv | CAGCAAGAGGAGTCCAAGTAAA |
| *krtt1c19e* | Fw | CACCACCAGCACCACTAAA |
| | Rv | CTTAACAAGGGACGTGGAAGAG |
| *lamb1a* | Fw | AGCCAGAAATCTGCTTCTACAG |
| | Rv | CTTTCTCCAGATCCACCAGTTC |
| *lamb2* | Fw | CATGCAGGACGCCAAGCAC |
| | Rv | CCTGGCTTTGCCTTCCAATACCTTC |
| *lamc2* | Fw | CAGCAGAGAGCCACCATTAC |
| | Rv | GGCCTTTCAATAGGTGGTATGT |
| *mcm6* | Fw | GGAGCTCATCAGTAAGAAGACC |
| | Rv | GGACTCAGCAGACCCTTTC |
| *mgst2* | Fw | TTTCGTGGTGCTATTGTGGATA |
| | Rv | GCCGGTAAAGTACATCTCTCTG |
| *mmp9* | Fw | AGAAGCTCGGCCTACCAAGCGAC |
| | Rv | GGGTATCCTCTGTCAATCAGCTGAGCC |
| *mmp13a* | Fw | CCTCAGAGCCCAGATGTTGAACTGC |
| | Rv | CTGTCTTCCTTGTAGAGGACTGCGTC |
| *opn1lw2* | Fw | ATCACCTGTTGCATCCTTCC |
| | Rv | ACACTTCCTTCTCGGCCTTC |
| *opn1mw1* | Fw | ACTTTTGCATACCCGTCACC |
| | Rv | TCACTTCCCTCTCAGCCTTC |
| *plin2* | Fw | GTGATGGACTACCTGCTCAAC |
| | Rv | GGGTCCTCTGTTTGCTACATT |
| *smc1a* | Fw | CCAGGCCATTGTTATCTCTCTC |

**Supplementary Table S1.**
Primers sequences used in RT-qPCR.

| Gene | Primer | Sequence (5’ to 3’) |
|---|---|---|
| | Rv | GATCAAAGGTCAGCACCTTACT |
| *tdh2* | Fw | CCAGTGCGGTTTGATGATTCTA |
| | Rv | ATGTCTGTGACCAGCTCTGA |
| *timp2b* | Fw | GCACACTGCGCATAAACCTGTGTGA |
| | Rv | GGGATGATGGGACACTGGACAAT |

Fw: forward.
Rv: reverse.

**Supplementary Table S2**

Functional classification of the 25 most up- and down-regulated genes in 6 dpf lethal *adamtsl4* KO larvae compared to wild type controls. The fold change for each gene's expression is shown in Fig. 10A-B.

| Functional Group | Gene Names | Number of Genes |
|---|---|---|
| **Down-regulated** | | |
| **Uncharacterized Function** | *LOC100334024, si:ch211-220i18.4, zgc:136930, LOC100334908, si:ch211-213i16.3, si:ch73-264i18.3, LOC101886621, zgc:194125, si:dkey-159a18.1, zgc:92590, si:dkey-14d8.7, si:dkey-14d8.6, sycn.2* | 13 |
| **Ion Transport and Channel Activity** | *kcnv2b, slc14a2, si:dkey-202l22.6* | 3 |
| **Metabolism and Transport** | *fut9d, zgc:194125* | 2 |
| **Immune Response** | *ctsbb, LOC101885188, LOC101882067* | 3 |
| **Lens and Eye Development** | *crygm2c, crygm2f, crygm2b* | 3 |
| **Cell Death and Apoptosis** | *xkr8.2* | 1 |
| **Up-regulated** | | |
| **ECM and Cell Adhesion** | *mmp9, mmp13a, timp2b, lamc2, cd44a, LOC100333484 (integrin alpha-2), fam212b, si:ch211-241e1.3, si:ch1073-110a20.1 (mfap4.5), si:zfos-2330d3.8 (mfap4.8), si:ch1073-15f12.3 (lamb3)* | 11 |
| **Inflammatory Response and Immune Function** | *cxcl8a, lect2l, il11a, il1b, si:dkey-8k3.2 (c4b), LOC103910140* | 6 |
| **Stress Response and Protein Folding** | *hsp70.3, unc45b, si:ch211-199o1.2 (hspa1b)* | 3 |
| **Developmental Signaling and Transcription Regulation** | *fosl1a, adam8b* | 2 |
| **Metabolism** | *Gcga* | 1 |
| **Cytoskeletal organization** | *plek2* | 1 |
| **Uncharacterized Function** | ***LOC101883708*** | 1 |

**Supplementary Table S3**

Functional classification of the 25 most up- and down-regulated genes in 6 dpf wild type-like *adamtsl4* KO larvae compared to wild type controls. The fold change for each gene's expression is shown in Fig. 10C-D.

| Functional Group | Genes | Number of Genes |
|---|---|---|
| **Downregulated** | | |
| **Transcription Factors** | *stat1b, fosab, nfil3-5, rorcb, egr2a* | 5 |
| **Signaling Pathways** | *rgs13, gprc6a, si:dkey-57c15.4, trpv6* | 4 |
| **Protein Modulation/Degradation** | *hspb9, asb15b* | 2 |
| **Ion Channels and Transporters** | *kcnj1a.2, trpv6* | 2 |
| **Uncharacterized** | *LOC103909567, zgc:112492, si:ch211-114l13.1, si:ch73-264i18.3, zgc:162969, zgc:112368, zgc:92745, LOC100535423, si:dkey-159a18.1, zgc:92590, LOC101886621, zgc:194125* | 12 |
| **Upregulated** | | |
| **Circadian Rhythm Regulation** | *nr1d1, per1a, per1b, ciarta* | 4 |
| **Muscle and Structural Proteins** | *fbxo40.2, cmya5, casq1a, ankrd33aa, rtn2b* | 5 |
| **Vision and Phototransduction** | *grk7a, arr3a, arr3b, rpe65a, lratb.2* | 5 |
| **Lipid and Retinoid Metabolism** | *dhrs7ca, alox5b.3, eevs* | 3 |
| **Protein Modification and Degradation** | *si:ch73-23l24.1, si:ch211-132b12.7, usp21, wdr76, capn3b* | 5 |
| **Uncharacterized** | *LOC110438318, si:ch211-103a14.5, si:ch211-161h7.5* | 3 |

**Supplementary Table S4**

Functional classification of the 25 most up- and down-regulated genes in eyes of adult adamtsl4 KO compared to wild type controls. The fold change for each gene's expression is shown in Fig. 10E-F.

| Functional Group | Genes | Number of Genes |
|---|---|---|
| **Downregulated** | | |
| **Interferon-stimulated response** | *ifit10, ifit8, gig2g, gig2l, mxb* | 5 |
| **Hypothetical/unknown proteins** | *LOC103909357, LOC100001846, LOC100149931, LOC101884392, LOC110437826, LOC108179089, LOC799227, LOC108190663, LOC103911026, LOC110437832, LOC100332323, LOC110439931, LOC101885896* | 13 |
| **Zebrafish-specific genes** | *si:ch211-66k16.28, si:ch211-153b23.5, si:dkey-211g8.8, si:ch211-238e22.2, si:ch211-201o1.1* | 5 |
| **Chromosome-associated** | *cenpo* | 1 |
| **Other zebrafish genes** | *zgc:136683* | 1 |
| **Upregulated** | | |
| **Unknown/Uncharacterized** | *si:ch73-236c18.5, LOC100150791, LOC571282, LOC110438047, LOC108190572, LOC108179116, LOC101885837, si:dkey-29j8.3, si:dkey-242k1.4, LOC100334861, LOC108181187, LOC103908938, LOC103908728, LOC103910620, LOC101885874, LOC108182924, LOC799889, LOC101882679* | 18 |
| Cilia structure and function/**Cell adhesion** | *pcdh1a4, ccdc39,* ***si:dkey-254e1.1*** *(**cfap70)*** | 3 |
| **Immunity** | *mhc2dab* | 1 |
| **Iron-Sulfur Cluster** | *rfesd* | 1 |
| **Cohesion Complex** | *smc1a* | 1 |
| **Miscellaneous** | *zgc:153426* | 1 |

**Supplementary Table S5**

Functional ShinyGO pathway enrichment for all DEGs in lethal adamtsl4 KO zebrafish larvae (6 dpf) compared to wild type controls. The table highlights the 20 most significant biological pathways, grouped into functional classes and ranked by false discovery rate (FDR).

| Term | Number of genes | Number of genes in background | Genes | FDR | Description |
|---|---|---|---|---|---|
| **Immune and Stress-Related Responses** | | | | | |
| GO:0006952 | 80 | 868 | *adam8a, f3b, tnfb, tnfrsf1a, cyba, fgb, acp5a, zgc:65788, unm_sa911, ptgs2a, spon2b, anxa1a, fga, c4, sgk1, LOC563884, tnfaip3, caspa, spon2a, dbi, ncf1, c3c, c3a, ptger2b, LOC798235, cebpb, zgc:110354, pglyrp5, ch25h, adam8b, hyal6, zgc:123218, cfb, si:ch73-86n18.1, irf8, lygl1, LOC567869, Nox1, ano6, duox, irg1l, c5, card9, ptx3a, cpla2, zgc:172053, zgc:110251, tnfrsf1b, irf7, stat3, LOC794786, samhd1, zgc:172260, lacc1, LOC564660, TNIP2, f2rl1.2, LOC799204, irf3, c3b, ripk3l, cybb, bcl3, fn1a, ENSDARP00000113672, c7b, PARP4, hyal2, c7a, ifitm1, cxcr3.2, mst1ra, rnasel3, serpinf2b, tlr5a, tlr5b, elf3, tap1, ptk2ba, ltb4r* | 6.00E-16 | Defense response |
| GO:0006950 | 150 | 2723 | *adam8a, f3b, myofl, lamb2, fstl1a, pdcd6, hsp70, tnfb, tnfrsf1a, gpx1b, fosl1a, cyba, hsp90ab1, fgb, acp5a, zgc:65788, unm_sa911, hsp90aa1.1, ptgs2a, spon2b, hspa4a, pkd2, tcap, anxa1a, mgst1.1, fga, hsp70l, itgb4, c4, sgk1, dspa, LOC563884, tnfaip3, hig1, caspa, spon2a, dbi, serpine2, ncf1, nccrp1, tacr1a, tph1b, c3c, c3a, ptger2b, fgg, phlda3, bag3, mcm4, LOC798235, klf2a, cebpb, zgc:110354, loxl2a, pglyrp5, ch25h, adam8b, Flna, hyal6, cryaa, zgc:123218, cfb, f5, hspd1, si:ch73-86n18.1, irf8, lygl1, bves, prdx1, tgfbr2a, sdc4, LOC567869, hspb6, Nox1, ano6, map3k8, duox, irg1l, c5, card9, ENSDARP00000088309, ptx3a, cpla2, pak6b, f9b, zgc:172053, zgc:110251, sesn2, tnfrsf1b, irf7, stat3, LOC794786, samhd1, zgc:172260, lacc1, hbegfa, LOC564660, TNIP2, f2rl1.2, atf3, ppp1r15b, LOC799204, slc38a3b, plekhn1, irf3, c3b, zgc:174006, hbegfb, ETS1, asns, clu, dhcr24, ripk3l, cd44a, cybb, bcl3, fn1a, rbm24b, mvp, lmna, ENSDARP00000113672, c7b, myh9b, PARP4, hyal2, myof, cdkn1a, c7a, capn3b, dnajb1b, trpv4, ifitm1, cxcr3.2, stx2a, gpx3, mst1ra, f13a1b, gpx7, rnasel3, serpinf2b, tlr5a, tlr5b, elf3, tap1, egln3, ptk2ba, LOC559569, pik3cb, ltb4r, si:dkey-79d12.5* | 1.36E-12 | Response to stress |
| GO:0009605 | 110 | 1865 | *Opn1mw1, adam8a, per3, lamb2, per1b, fstl1a, gpr183, tnfb, epha2, fosl1a, cyba, bin2b, acsl4l, cry5, fgb, cmtm3, acp5a, unm_sa911, spon2b, lamc1, pkd2, tcap, rhogb, anxa1a, mgst1.1, fga, c4, cd63, LOC563884, hig1, caspa, spon2a, lect2l, LOC555409, serpine2, ncf1, c3c, c3a, ptger2b, bag3, LOC798235, cyp51, cebpb, zgc:110354, pglyrp5, opn1lw2, opn1lw1, ch25h, adam8b, irak3, hyal6, cfb, si:ch73-86n18.1, irf8, lygl1, LOC567869, ano6, irg1l, c5, card9, ptx3a, per1a, zgc:172053, itga6b, zgc:110251, sesn2, tnfrsf1b, irf7, LOC794786, samhd1, zgc:172260, lacc1, hbegfa, MYOT, LOC564660, TNIP2, akap12b, f2rl1.2, atf3, slc38a3b, irf3, c3b, LOC793246, hbegfb, ETS1, junba, asns, ripk3l, cybb, plxnd1, bcl3, adora2b, LOC564958, ENSDARP00000113672, si:ch1073-110a20.1, c7b, hyal2, cdkn1a, c7a, si:ch73-63e15.2, trpv4, ifitm1, cxcr3.2, mst1ra, met, rnasel3, tlr5a, tlr5b, ptk2ba, si:dkey-79d12.5* | 3.16E-10 | Response to external stimulus |

**Supplementary Table S5**

Functional ShinyGO pathway enrichment for all DEGs in lethal adamtsl4 KO zebrafish larvae (6 dpf) compared to wild type controls. The table highlights the 20 most significant biological pathways, grouped into functional classes and ranked by false discovery rate (FDR).

| Term | Number of genes | Number of genes in background | Genes | FDR | Description |
|---|---|---|---|---|---|
| GO:0006954 | 39 | 339 | *adam8a, f3b, tnfrsf1a, cyba, zgc:65788, ptgs2a, anxa1a, c4, sgk1, LOC563884, tnfaip3, c3c, c3a, ptger2b, LOC798235, cebpb, adam8b, hyal6, zgc:123218, ano6, irg1l, c5, cpla2, zgc:110251, tnfrsf1b, stat3, lacc1, TNIP2, f2rl1.2, LOC799204, c3b, cybb, fn1a, PARP4, serpinf2b, tlr5a, tlr5b, elf3, ltb4r* | 1.51E-09 | Inflammatory response |
| GO:0009607 | 65 | 906 | *tnfb, cyba, fgb, acp5a, unm_sa911, spon2b, anxa1a, mgst1.1, fga, c4, LOC563884, caspa, spon2a, LOC555409, ncf1, c3c, c3a, ptger2b, LOC798235, cyp51, cebpb, zgc:110354, pglyrp5, ch25h, irak3, hyal6, cfb, si:ch73-86n18.1, irf8, lygl1, LOC567869, irg1l, c5, card9, ptx3a, zgc:172053, tnfrsf1b, irf7, LOC794786, samhd1, zgc:172260, lacc1, LOC564660, TNIP2, akap12b, f2rl1.2, ppp1r15b, irf3, c3b, junba, ripk3l, cybb, bcl3, LOC564958, ENSDARP00000113672, c7b, hyal2, c7a, si:ch73-63e15.2, ifitm1, cxcr3.2, mst1ra, rnasel3, tlr5a, tlr5b* | 2.02E-08 | Response to biotic stimulus |
| GO:0051707 | 64 | 884 | *tnfb, cyba, fgb, acp5a, unm_sa911, spon2b, anxa1a, mgst1.1, fga, c4, LOC563884, caspa, spon2a, LOC555409, ncf1, c3c, c3a, ptger2b, LOC798235, cyp51, cebpb, zgc:110354, pglyrp5, ch25h, irak3, hyal6, cfb, si:ch73-86n18.1, irf8, lygl1, LOC567869, irg1l, c5, card9, ptx3a, zgc:172053, tnfrsf1b, irf7, LOC794786, samhd1, zgc:172260, lacc1, LOC564660, TNIP2, akap12b, f2rl1.2, irf3, c3b, junba, ripk3l, cybb, bcl3, LOC564958, ENSDARP00000113672, c7b, hyal2, c7a, si:ch73-63e15.2, ifitm1, cxcr3.2, mst1ra, rnasel3, tlr5a, tlr5b* | 2.02E-08 | Response to other organism |
| GO:0098542 | 50 | 614 | *tnfb, cyba, fgb, acp5a, unm_sa911, spon2b, anxa1a, fga, c4, caspa, spon2a, ncf1, c3c, c3a, cebpb, zgc:110354, pglyrp5, ch25h, cfb, si:ch73-86n18.1, irf8, lygl1, LOC567869, irg1l, c5, card9, ptx3a, zgc:172053, irf7, LOC794786, samhd1, zgc:172260, lacc1, LOC564660, f2rl1.2, irf3, c3b, ripk3l, cybb, bcl3, ENSDARP00000113672, c7b, hyal2, c7a, ifitm1, cxcr3.2, mst1ra, rnasel3, tlr5a, tlr5b* | 6.15E-08 | Defense response to other organism |
| GO:0001819 | 33 | 299 | *adam8a, trim16, tnfb, cyba, spon2b, hhla2a.2, anxa1a, LOC563884, spon2a, cebpb, adam8b, LOC557072, hspd1, irf8, LOC567869, Nox1, card9, zgc:172053, irf7, casp8, akap12b, f2rl1.2, LOC799204, irf3, Runx1, bcl3, adora2b, hyal2, trpv4, cd276, serpinf2b, tlr5a, tlr5b* | 1.12E-07 | Positive regulation of cytokine production |
| GO:0050727 | 28 | 219 | *adam8a, per1b, tnfb, tnfrsf1a, acp5a, anxa1a, tnfaip3, dnase1l3l, cebpa, LOC798235, cebpb, pglyrp5, adam8b, LOC567869, tnfaip6, irg1l, per1a, BCL6B, zgc:110251, lacc1, nr1d1, LOC799204, ETS1, nr1d2b, adora2b, hyal2, si:ch73-63e15.2, trpv4* | 1.17E-07 | Regulation of inflammatory response |
| GO:0070555 | 16 | 71 | *cyba, LOC798235, klf2a, cebpb, irak3, hyal6, LOC567869, irg1l, ADAMTS12, ENSDARP00000088309, TNIP2, akap12b, nr1d1, LOC799204, ETS1, hyal2* | 7.03E-07 | Response to interleukin-1 |
| **Developmental Processes** | | | | | |

**Supplementary Table S5**

Functional ShinyGO pathway enrichment for all DEGs in lethal adamtsl4 KO zebrafish larvae (6 dpf) compared to wild type controls. The table highlights the 20 most significant biological pathways, grouped into functional classes and ranked by false discovery rate (FDR).

| Term | Number of genes | Number of genes in background | Genes | FDR | Description |
|---|---|---|---|---|---|
| GO:0001654 | 40 | 429 | *lamb2, inhbaa, epha2, unc45b, lamc1, cryba2a, cryba4, zgc:136220, crygm5, LOC569604, cryaa, bves, tgfbr2a, si:dkey-57a22.15, crygm2d6, crygm2d5, crygm2d12, crygm2d8, crygm2d15, crygm2d11, stat3, crygmx, Crygm2d4, crygm2d16, crygm2d7, crygm2d9, crygm2d20, zgc:171792, tmod1, LOC565849, crygm2d13, crygm2d10, crygm2d2, mvp, crygm2d3, crygm2d1, cryba1a, crybb1, frem3, crygm2d21* | 1.17E-07 | Eye development |
| GO:0048880 | 41 | 460 | *lamb2, inhbaa, epha2, unc45b, lamc1, neurod, cryba2a, cryba4, zgc:136220, crygm5, LOC569604, cryaa, bves, tgfbr2a, si:dkey-57a22.15, crygm2d6, crygm2d5, crygm2d12, crygm2d8, crygm2d15, crygm2d11, stat3, crygmx, Crygm2d4, crygm2d16, crygm2d7, crygm2d9, crygm2d20, zgc:171792, tmod1, LOC565849, crygm2d13, crygm2d10, crygm2d2, mvp, crygm2d3, crygm2d1, cryba1a, crybb1, frem3, crygm2d21* | 1.85E-07 | Sensory system development |
| GO:0048513 | 144 | 3102 | *nradd, cldn7b, lamb2, inhbaa, pdcd6, gpr183, tnfb, epha2, fosl1a, klhl40b, rras, hsp90ab1, acp5a, hsp90aa1.1, fads2, unc45b, lamc1, neurod, pkd2, tcap, itgb4, dspa, LOC563884, zgc:153629, caspa, serpinh1b, cryba2a, flvcr1, serpine2, cryba4, zgc:136220, arrdc3b, cebpa, tph1b, acta1a, b4galt5, dvr1, bag3, fgf8b, acta1b, nab1b, bmp1b, crygm5, mthfd1l, Si:ch73-187m15.4, klf2a, cebpb, fgf1b, srd5a2a, loxl2a, LOC569604, krtt1c19e, ENSDARP00000067092, Flna, hyal6, LOC557072, cryaa, ankrd33aa, irf8, murcb, zgc:86709, bves, tgfbr2a, si:ch1073-15f12.3, slc25a38a, sdc4, LOC567869, ano6, si:dkey-57a22.15, LAMC2, slc25a37, cpla2, crygm2d6, crygm2d5, crygm2d12, crygm2d8, crygm2d15, crygm2d11, itga6b, gfer, tnfrsf1b, stat3, casp8, crygmx, meox1, ENSDARP00000096826, Crygm2d4, snx10a, iqgap1, crygm2d16, SLC4A7, wnt4b, crygm2d7, crygm2d9, crygm2d20, zgc:171792, tmod1, zgc:175175, LOC565849, cd248b, f2rl1.2, atf3, crygm2d13, slc38a3b, crygm2d10, ETS1, junba, mustn1, dhcr24, ripk3l, crygm2d2, c1galt1b, Runx1, plxnd1, bcl3, fosl2, rbm24b, mvp, crygm2d3, lmna, crygm2d1, rap1b, myh9b, cthrc1a, hyal2, cdkn1a, ptpn3, LOC567640, cryba1a, si:ch73-63e15.2, trpv4, crybb1, frem3, actn1, crygm2d21, xirp2a, actl6b, elf3, LOC562247, ptk2ba, PLCE1, synpo2lb, col12a1a, si:dkey-79d12.5* | 2.22E-07 | Animal organ development |
| GO:0043010 | 34 | 349 | *lamb2, inhbaa, epha2, unc45b, cryba2a, cryba4, zgc:136220, crygm5, LOC569604, cryaa, bves, si:dkey-57a22.15, crygm2d6, crygm2d5, crygm2d12, crygm2d8, crygm2d15, crygm2d11, crygmx, Crygm2d4, crygm2d16, crygm2d7, crygm2d9, crygm2d20, zgc:171792, tmod1, crygm2d13, crygm2d10, crygm2d2, crygm2d3, crygm2d1, cryba1a, crybb1, crygm2d21* | 7.03E-07 | Camera-type eye development |
| GO:0007423 | 48 | 636 | *cldn7b, lamb2, inhbaa, epha2, unc45b, lamc1, neurod, tcap, cryba2a, cryba4, zgc:136220, cebpa, b4galt5, crygm5, LOC569604, cryaa, bves, tgfbr2a, sdc4, si:dkey-57a22.15, crygm2d6, crygm2d5, crygm2d12, crygm2d8, crygm2d15, crygm2d11, stat3, crygmx, Crygm2d4, crygm2d16, SLC4A7, crygm2d7, crygm2d9, crygm2d20, zgc:171792, tmod1,* | 7.48E-07 | Sensory organ development |

**Supplementary Table S5**

Functional ShinyGO pathway enrichment for all DEGs in lethal adamtsl4 KO zebrafish larvae (6 dpf) compared to wild type controls. The table highlights the 20 most significant biological pathways, grouped into functional classes and ranked by false discovery rate (FDR).

| Term | Number of genes | Number of genes in background | Genes | FDR | Description |
|---|---|---|---|---|---|
| | | | *LOC565849, crygm2d13, crygm2d10, crygm2d2, mvp, crygm2d3, crygm2d1, cthrc1a, cryba1a, crybb1, frem3, crygm2d21* | | |

**Supplementary Table S6**

Functional ShinyGO pathway enrichment of all DEGs in wild type-like adamtsl4 KO zebrafish larvae (6 dpf) compared to wild type controls. The table highlights the 20 most significant biological pathways, grouped into functional classes and ranked by false discovery rate (FDR).

| Term | Number of genes | Number of genes in background | Genes | FDR | Description |
|---|---|---|---|---|---|
| **Circadian Rhythm and Rhythmic Processes** | | | | | |
| GO:0048511 | 11 | 267 | *per1b, cry5, ciart, tefa, per1a, nr1d1, rorcb, nr1d2b, si:ch211-132b12.7, nfil3-5, cipca* | 5.72E-05 | Rhythmic process |
| GO:0042752 | 7 | 121 | *per1b, cry5, per1a, nr1d1, nr1d2b, si:ch211-132b12.7, cipca* | 2.20E-03 | Regulation of circadian rhythm |
| GO:0007623 | 7 | 146 | *per1b, cry5, ciart, per1a, nr1d1, nr1d2b, nfil3-5* | 5.00E-03 | Circadian rhythm |
| GO:0042754 | 3 | 16 | *cry5, si:ch211-132b12.7, cipca* | 4.02E-02 | Negative regulation of circadian rhythm |
| **Visual and Light Response** | | | | | |
| GO:0007601 | 7 | 188 | *rpe65a, pdca, zgc:153443, grk7a, zgc:73075, rp1l1b, cnga3a* | 1.85E-02 | Visual perception |
| GO:0009416 | 7 | 258 | *per1b, cry5, xpc, zgc:173915, grk7a, per1a, zgc:73075* | 4.09E-02 | Response to light stimulus |
| **DNA Replication and Repair** | | | | | |
| GO:0006267 | 3 | 10 | *mcm3, mcm5, mcm6* | 2.21E-02 | Pre-replicative complex assembly involved in nuclear cell cycle DNA replication |
| GO:0000727 | 3 | 14 | *mcm3, mcm5, mcm6* | 3.12E-02 | Double-strand break repair via break-induced replication |
| **Signaling Pathways and Regulation** | | | | | |
| GO:0043124 | 4 | 42 | *per1b, per1a, nr1d1, stat1b* | 2.69E-02 | Negative regulation of I-kappaB kinase/NF-kappaB signaling |
| GO:2000322 | 3 | 16 | *per1b, cry5, per1a* | 4.02E-02 | Regulation of glucocorticoid receptor signaling pathway |

**Supplementary Table S7**

Functional ShinyGO pathway enrichment of all DEGs in eyes of adult adamtsl4 KO zebrafish compared to wild type controls. The table highlights the 20 most significant biological pathways, grouped into functional classes and ranked by false discovery rate (FDR).

| Term | Number of genes | Number of genes in background | Genes | FDR | Description |
|---|---|---|---|---|---|
| **Cell adhesion and extracellular matrix** | | | | | |
| GO:0007155 | 79 | 697 | *si:dkey-32n7.4, obsl1a, lmo7a, ncanb, robo4, cldnb, cldn5b, ctnnal1, itga2b, thbs4b, dspa, si:ch211-241e1.3, ptk2ba, pkp2, si:ch211-106h11.3, itgb8, podxl, lamb2l, plxnb2b, si:dkey-24p1.1, dsc2l, si:dkey-15j16.3, cldne, cldn5a, spp1, cldn19, pkp3a, cdh27, clstn2, CLSTN2, si:ch73-233f7.1, vcam1b, lyve1b, gpnmb, zgc:153759, zgc:153631, si:ch211-203k16.3, frem1a, zgc:136892, cldna, jupa, itga1, si:ch73-380l3.2, pcdh12, tnfrsf18, si:ch211-66e2.3, ninj2, si:ch211-214p13.3, cd44a, igsf5a, pcdh2aa15, macf1b, ncam3, pcdh2ab9, si:dkey-22i16.9, itgam, si:ch73-181m17.1, pcdh1a4, si:dkey-238d18.7, si:ch211-39f2.3, si:ch211-149e23.4, itgae.1, si:ch73-329n5.6, comp, cdhr5b, zmp:0000001082, si:ch73-379j16.2, pcdh1g3, pcdh2ac, anxa1b, crb1, lamb1a, ntn4, si:ch73-233f7.3, mag, postnb, anxa1c, cxcl8a, lurap1* | 3.26E-04 | Cell adhesion |
| GO:0022610 | 79 | 697 | *si:dkey-32n7.4, obsl1a, lmo7a, ncanb, robo4, cldnb, cldn5b, ctnnal1, itga2b, thbs4b, dspa, si:ch211-241e1.3, ptk2ba, pkp2, si:ch211-106h11.3, itgb8, podxl, lamb2l, plxnb2b, si:dkey-24p1.1, dsc2l, si:dkey-15j16.3, cldne, cldn5a, spp1, cldn19, pkp3a, cdh27, clstn2, CLSTN2, si:ch73-233f7.1, vcam1b, lyve1b, gpnmb, zgc:153759, zgc:153631, si:ch211-203k16.3, frem1a, zgc:136892, cldna, jupa, itga1, si:ch73-380l3.2, pcdh12, tnfrsf18, si:ch211-66e2.3, ninj2, si:ch211-214p13.3, cd44a, igsf5a, pcdh2aa15, macf1b, ncam3, pcdh2ab9, si:dkey-22i16.9, itgam, si:ch73-181m17.1, pcdh1a4, si:dkey-238d18.7, si:ch211-39f2.3, si:ch211-149e23.4, itgae.1, si:ch73-329n5.6, comp, cdhr5b, zmp:0000001082, si:ch73-379j16.2, pcdh1g3, pcdh2ac, anxa1b, crb1, lamb1a, ntn4, si:ch73-233f7.3, mag, postnb, anxa1c, cxcl8a, lurap1* | 3.26E-04 | Biological adhesion |
| GO:0018149 | 8 | 15 | *tgm1, f13a1a.1, ndnf, tgm2l, tgm8, tgm5l, tgm1l4, tgm1l1* | 1.11E-03 | Peptide cross-linking |
| **Developmental Processes** | | | | | |
| GO:0009653 | 142 | 1630 | *meis3, meis1a, sox9a, obsl1a, marcksa, tbx21, grna, odc1, sema3bl, foxl1, dbnla, smyd1a, robo4, foxi3b, cacnb4a, calcrlb, rgma, col11a2, sema3gb, wnt11, acvrl1, itga2b, dusp5, sparc, sec23b, sp7, tnnt2a, nkx6.1, si:ch211-241e1.3, ptk2ba, pkp2, kcnh1b, socs3a, gmds, extl3, palm1a, igf1ra, itgb8, ahsa1a, stat4, tnni2a.4, foxd1, sox7, hspa12b, foxq1a, fgf10a, ccdc80l1, podxl, pacsin1a, lamb2l, angptl4, ntn5, plxnb2b, socs1a, pkdccb, pigp, wnt3, fli1b, prickle1a, si:dkey-49n23.1, ppih, actc1a, klf2a, cldne, atg4da, rab3ab, sox9b, slit1a, chad, dkk1b, rbpms2a, inab, csf1rb, pax9, etv2, ostn, slc8a4a, snai1a, cxcr4a, ephb6, foxd2, draxin, sox18, tax1bp3, si:ch1073-15f12.3, plod1a, kcnb1, vcam1b, ndnf, dagla, lamc2, vgll4l, otog, dmrt2b, jupa, fhl1a, crabp2a, gas7a, birc5a, sema5bb, pigt, tbr1a, si:ch211-159i8.4, ptprb, frem2a, sox10, f2rl1.2, sox11a, grhl3, sema7a, foxe1, btbd3b, si:dkey-1c11.1, smoc1, ncam3, si:dkey-6b12.5, myh6, smyd1b, cyt1, nyap2b, rac1l, efna3b, iqgap3, pygo1, rgs2, anxa1b, pbx1a, crb1, lamb1a, irx1a, irf6, ntn4, ved, hmgcs1, clec14a, spry1, igfbp7, anxa1c, gipc1, cxcl8a, fgf4, lurap1* | 2.32E-02 | Anatomical structure morphogenesis |

**Supplementary Table S7**
Functional ShinyGO pathway enrichment of all DEGs in eyes of adult adamtsl4 KO zebrafish compared to wild type controls. The table highlights the 20 most significant biological pathways, grouped into functional classes and ranked by false discovery rate (FDR).

| Term | Number of genes | Number of genes in background | Genes | FDR | Description |
|---|---|---|---|---|---|
| GO:0048513 | 158 | 1887 | *meis3, meis1a, loxa, sox9a, obsl1a, marcksa, tbx21, grna, pou4f1, odc1, sema3bl, foxl1, crygm4, myo1eb, smyd1a, cldnb, clocka, col11a2, pygmb, tnfb, sema3gb, dkk1a, myl1, otpa, npm1a, wnt11, klf3, klf1, acvrl1, heg1, sparc, sec23b, sp7, tnnt2a, klhl24a, nkx6.1, si:ch211-241e1.3, ptk2ba, pkp2, kcnh1b, fzd5, socs3a, gmds, extl3, igf1ra, adcyap1b, ahsa1a, sox7, fgf10a, podxl, csrnp1a, lamb2l, irx5a, xbp1, irx4a, ntn5, slc43a1a, socs1a, pkdccb, wnt3, ca8, mpl, fstl1b, prickle1a, si:dkey-49n23.1, pklr, actc1a, klf2a, cx30.3, tyms, atg4da, cldn5a, sox9b, spp1, dkk1b, mybpc1, irf7, rtn4rl2a, rag1, tnni1b, hmgcra, rtkn2a, csf1rb, myb, pax9, crip1, etv2, ostn, armc4, slc8a4a, rel, hsp70l, irx1b, tmem88a, snai1a, cxcr4a, epas1b, six2a, draxin, si:ch1073-15f12.3, kcnb1, haus3, ndnf, tmem119b, klf9, lamc2, vgll4l, otog, zgc:153115, pou4f2, jupa, fhl1a, crabp2a, birc5a, sema5bb, shox2, sez6l2, stat1b, tbr1a, pou3f2b, cdkn1a, sox10, igsf10, sox11a, grhl3, sema7a, foxe1, gcga, mgp, ifnlr1, smoc1, clpb, myh6, smyd1b, rbp7a, dio2, tmem119a, pygo1, rgs2, elovl1a, anxa1b, pbx1a, zfyve9b, hif1al, lamb1a, irx1a, irf6, ntn4, ved, crb3a, hmgcs1, spry1, postnb, anxa1c, cxcl8a, fgf4, si:ch211-243a20.4, lurap1* | 3.15E-02 | Animal organ development |
| GO:0009887 | 58 | 559 | *meis3, meis1a, sox9a, obsl1a, marcksa, tbx21, odc1, col11a2, wnt11, sparc, sec23b, sp7, tnnt2a, si:ch211-241e1.3, pkp2, socs3a, gmds, igf1ra, ahsa1a, fgf10a, lamb2l, ntn5, socs1a, pkdccb, prickle1a, actc1a, klf2a, sox9b, csf1rb, pax9, etv2, ostn, slc8a4a, cxcr4a, si:ch1073-15f12.3, ndnf, lamc2, vgll4l, otog, fhl1a, tbr1a, sox10, sox11a, grhl3, foxe1, smoc1, myh6, pygo1, pbx1a, lamb1a, irx1a, irf6, ntn4, ved, hmgcs1, spry1, fgf4, lurap1* | 3.69E-02 | Animal organ morphogenesis |
| GO:0033688 | 3 | 3 | *sp7, tmem119b, tmem119a* | 3.98E-02 | Regulation of osteoblast proliferation |
| GO:0048731 | 216 | 2754 | *meis3, meis1a, loxa, sox9a, igf2bp2a, obsl1a, marcksa, tbx21, grna, pou4f1, ncanb, odc1, sema3bl, foxl1, dbnla, crygm4, myo1eb, smyd1a, robo4, cldnb, calcrlb, clocka, plp1b, rgma, col11a2, pygmb, tnfb, sema3gb, dkk1a, myl1, otpa, npm1a, wnt11, klf3, klf1, acvrl1, heg1, itga2b, sparc, sec23b, sp7, tnnt2a, klhl24a, nkx6.1, si:ch211-241e1.3, ptk2ba, pkp2, kcnh1b, fzd5, socs3a, gmds, extl3, igf1ra, adcyap1b, itgb8, ahsa1a, stat4, sox7, hspa12b, fgf10a, ccdc80l1, podxl, csrnp1a, pacsin1a, lamb2l, irx5a, xbp1, irx4a, angptl4, ntn5, plxnb2b, slc43a1a, socs1a, pkdccb, stmn3, wnt3, ca8, mpl, fstl1b, neurod6a, fli1b, prickle1a, si:dkey-49n23.1, ppih, pklr, actc1a, klf2a, cx30.3, tyms, zgc:153665, atg4da, cldn5a, rab3ab, sox9b, spp1, slit1a, chad, dkk1b, gsnb, mybpc1, irf7, rbpms2a, rtn4rl2a, rag1, tnni1b, hmgcra, inab, rtkn2a, csf1rb, myb, pax9, crip1, etv2, ostn, armc4, slc8a4a, rel, foxc1b, hsp70l, irx1b, tmem88a, snai1a, cxcr4a, epas1b, ephb6, six2a, draxin, sox18, nav1b, si:ch1073-15f12.3, kcnb1, clstn2, atxn1b, haus3, si:ch73-233f7.1, vcam1b, ndnf, dagla, manf, tmem119b, klf9, lamc2, vgll4l, otog, zgc:153115, pou4f2, dmrt2b, jupa, fhl1a, nrn1a, crabp2a,* | 3.98E-02 | System development |

**Supplementary Table S7**

Functional ShinyGO pathway enrichment of all DEGs in eyes of adult adamtsl4 KO zebrafish compared to wild type controls. The table highlights the 20 most significant biological pathways, grouped into functional classes and ranked by false discovery rate (FDR).

| Term | Number of genes | Number of genes in background | Genes | FDR | Description |
|---|---|---|---|---|---|
| | | | *rbfox3b, gas7a, birc5a, sema5bb, shox2, sez6l2, stat1b, tbr1a, pou3f2b, si:ch211-159i8.4, cdkn1a, ptprb, sox10, igsf10, sox11a, lmx1al, gal3st1b, grhl3, myrf, sema7a, foxe1, gcga, mgp, ifnlr1, btbd3b, si:dkey-1c11.1, smoc1, ncam3, clpb, myh6, smyd1b, foxc1a, rbp7a, nyap2b, dio2, tmem119a, si:dkey-221l4.10, efna3b, pygo1, rgs2, pclob, pcdh2ac, elovl1a, anxa1b, pbx1a, zfyve9b, hif1al, lamb1a, irx1a, etv1, irf6, ntn4, ved, crb3a, hmgcs1, clec14a, spry1, igfbp7, postnb, anxa1c, gipc1, cxcl8a, fgf4, si:ch211-243a20.4, lurap1* | | |
| **Metabolism Processes** | | | | | |
| GO:0005978 | 8 | 27 | *gys2, zgc:77112, gys1, gbe1a, phkg1b, ppp1r3ca, pask, ppp1r3da* | 3.98E-02 | Glycogen biosynthetic process |
| GO:0009250 | 8 | 27 | *gys2, zgc:77112, gys1, gbe1a, phkg1b, ppp1r3ca, pask, ppp1r3da* | 3.98E-02 | Glucan biosynthetic process |
| **Immune and Stress-Related Responses** | | | | | |
| GO:0002376 | 104 | 1141 | *meis1a, tbx21, ifit10, clocka, c3a.1, tnfb, npm1a, klf3, klf1, ptk6a, sparc, sec23b, si:ch1073-280e3.1, tnnt2a, ptk2ba, coch, extl3, nos2a, irf10, fgf10a, csrnp1a, tnfsf10l3, ccl19a.2, tlr2, socs1a, zgc:171687, MFAP4, ccr6b, mpl, ccl19b, tlr18, si:dkey-15j16.3, fpr1, mpeg1.2, irf1b, psmb10, cd226, irf7, saa, rag1, pkz, gpatch3, zgc:153932, rtkn2a, csf1rb, myb, etv2, rel, hsp70l, foxp3a, tmem88a, tnfsf10, cxcr4a, epas1b, tapbpl, ccl19a.1, themis2, rxfp3.3b, zgc:153654, haus3, klf9, tlr7, gbp4, zgc:153759, zgc:153631, rxfp3.3a1, rxfp3.3a3, cd4-1, zmp:0000001138, cxcl32b.1, ctss2.1, ccl44, si:dkey-79f11.5, si:ch73-211l2.3, stat1b, cxcr3.1, sema7a, ackr4a, mhc2dab, dhx58, socs1b, si:ch211-66k16.27, si:dkey-193b15.5, cd180, ccl34b.4, si:dkey-192g7.3, ccl20b, si:dkey-220k22.3, si:ch211-197g15.8, si:dkey-22i16.9, si:dkey-117a8.1, anxa1b, pbx1a, tlr1, zgc:123107, irf6, cxcl8b.1, hmgcs1, si:rp71-81e14.2, dicp1.17, anxa1c, leap2, cxcl8a, si:ch211-243a20.4* | 2.90E-02 | Immune system process |
| GO:0006955 | 71 | 725 | *ifit10, c3a.1, tnfb, ptk6a, si:ch1073-280e3.1, ptk2ba, coch, nos2a, tnfsf10l3, ccl19a.2, tlr2, socs1a, zgc:171687, MFAP4, ccr6b, ccl19b, tlr18, si:dkey-15j16.3, fpr1, mpeg1.2, irf1b, cd226, saa, pkz, gpatch3, rel, tnfsf10, cxcr4a, ccl19a.1, themis2, rxfp3.3b, zgc:153654, tlr7, gbp4, zgc:153759, zgc:153631, rxfp3.3a1, rxfp3.3a3, cd4-1, zmp:0000001138, cxcl32b.1, ctss2.1, ccl44, si:dkey-79f11.5, si:ch73-211l2.3, stat1b, cxcr3.1, sema7a, ackr4a, mhc2dab, dhx58, socs1b, si:ch211-66k16.27, si:dkey-193b15.5, cd180, ccl34b.4, si:dkey-192g7.3, ccl20b, si:dkey-220k22.3, si:ch211-197g15.8,* | 3.69E-02 | Immune response |

**Supplementary Table S7**

Functional ShinyGO pathway enrichment of all DEGs in eyes of adult adamtsl4 KO zebrafish compared to wild type controls. The table highlights the 20 most significant biological pathways, grouped into functional classes and ranked by false discovery rate (FDR).

| Term | Number of genes | Number of genes in background | Genes | FDR | Description |
|---|---|---|---|---|---|
| | | | *si:dkey-22i16.9, si:dkey-117a8.1, anxa1b, tlr1, zgc:123107, cxcl8b.1, si:rp71-81e14.2, anxa1c, leap2, cxcl8a, si:ch211-243a20.4* | | |
| GO:0009607 | 47 | 440 | *nitr2b, ifit10, zgc:77112, ptk6a, mpx, si:ch1073-280e3.1, ptk2ba, coch, nos2a, ldlra, ccl19a.2, angptl4, tlr2, socs1a, zgc:171687, MFAP4, mcm4, tlr18, si:dkey-15j16.3, mpeg1.2, irf1b, cd226, irf7, saa, pkz, gpatch3, rel, f5, ccl19a.1, irg1l, gbp4, zgc:153759, zgc:153631, cxcl32b.1, ccl44, stat1b, dhx58, socs1b, si:ch211-66k16.27, cd180, ccl20b, si:dkey-22i16.9, nsdhl, tlr1, cxcl8b.1, leap2, cxcl8a* | 3.98E-02 | Response to biotic stimulus |
| GO:0031347 | 13 | 67 | *SBNO2, coch, tlr2, socs1a, irf1b, cd226, gpatch3, sema7a, socs1b, anxa1b, tlr1, anxa1c, cxcl8a* | 3.98E-02 | Regulation of defense response |
| GO:0034097 | 30 | 237 | *ghrb, crlf1a, stat4, ccl19a.2, socs1a, ccr6b, il12rb2, csf1rb, rel, cxcr4a, ccl19a.1, rxfp3.3b, rxfp3.3a1, rxfp3.3a3, cd4-1, cxcl32b.1, cntf, ccl44, stat1b, tnfrsf18, cxcr3.1, ackr4a, ifnlr1, socs1b, ccl20b, anxa1b, cxcl8b.1, dicp1.17, anxa1c, cxcl8a* | 3.98E-02 | Response to cytokine |
| GO:0043207 | 47 | 438 | *nitr2b, ifit10, zgc:77112, ptk6a, mpx, si:ch1073-280e3.1, ptk2ba, coch, nos2a, ldlra, ccl19a.2, angptl4, tlr2, socs1a, zgc:171687, MFAP4, mcm4, tlr18, si:dkey-15j16.3, mpeg1.2, irf1b, cd226, irf7, saa, pkz, gpatch3, rel, f5, ccl19a.1, irg1l, gbp4, zgc:153759, zgc:153631, cxcl32b.1, ccl44, stat1b, dhx58, socs1b, si:ch211-66k16.27, cd180, ccl20b, si:dkey-22i16.9, nsdhl, tlr1, cxcl8b.1, leap2, cxcl8a* | 3.98E-02 | Response to external biotic stimulus |
| GO:0042330 | 44 | 404 | *sema3bl, smyd1a, robo4, sema3gb, si:ch211-241e1.3, ptk2ba, extl3, fgf10a, ccdc80l1, ccl19a.2, ntn5, plxnb2b, ccr6b, si:dkey-49n23.1, ccr7, slit1a, chad, rbpms2a, saa, cxcr4a, ephb6, draxin, ccl19a.1, rxfp3.3b, si:ch1073-15f12.3, rxfp3.3a1, rxfp3.3a3, pdgfra, cxcl32b.1, ccl44, sema5bb, cxcr3.1, sema7a, ackr4a, si:dkey-1c11.1, ncam3, clpb, ccl20b, efna3b, anxa1b, lamb1a, cxcl8b.1, anxa1c, cxcl8a* | 3.98E-02 | Taxis |
| **Cellular architecture** | | | | | |
| GO:0045103 | 8 | 23 | *evpla, nefma, dspa, pkp2, nefmb, macf1b, eppk1, ppl* | 2.69E-02 | Intermediate filament-based process |
| GO:0045104 | 8 | 23 | *evpla, nefma, dspa, pkp2, nefmb, macf1b, eppk1, ppl* | 2.69E-02 | Intermediate filament cytoskeleton organization |

**Supplementary Table S7**

Functional ShinyGO pathway enrichment of all DEGs in eyes of adult adamtsl4 KO zebrafish compared to wild type controls. The table highlights the 20 most significant biological pathways, grouped into functional classes and ranked by false discovery rate (FDR).

| Term | Number of genes | Number of genes in background | Genes | FDR | Description |
|---|---|---|---|---|---|
| GO:0045110 | 3 | 3 | *nefma, pkp2, nefmb* | 3.98E-02 | Intermediate filament bundle assembly |
